# Z-DNA Reveals a Conformational Dimension of Neutrophil Extracellular Trap Organization

**DOI:** 10.64898/2026.05.04.722750

**Authors:** Tevriz Dilan Demir, Nikhil Ram-Mohan, Samuel Yang

## Abstract

Neutrophil extracellular traps (NETs), with their networked extracellular DNA (eDNA), are key effectors of innate immunity which when ineffectively cleared can lead to immunopathologies. Here, we demonstrate intrinsic, host-derived left-handed noncanonical Z-DNA in NETs eDNA. Immunofluorescence imaging revealed Z-DNA forming extensive filaments tracing the non-Z-DNA backbone across diverse NETosis stimuli, while circular dichroism independently supported its presence. Temporal analysis revealed an early mitochondria-associated component with preferential ZBP1 association, followed by progressive accumulation within the eDNA network as NETs matured. Z-DNA accumulation was associated with progressive changes in NET architecture and topology, including network organization, branching, junctional complexity, filament thickness, and pore organization. Chemical perturbation of the Z:non-Z equilibrium was accompanied by NET architectural remodeling, associating DNA conformational state with higher-order scaffold organization. Z-DNA showed greater relative retention than non-Z-DNA following nuclease treatment, whereas chloroquine-mediated shifting toward a non-Z state reduced this preferential retention. Finally, Z-DNA–enriched NET fractions induced greater IFN-α production by human PBMCs than non-Z-DNA fractions. These findings identify Z-DNA as a dynamic component of NETs whose abundance and spatial organization are associated with NET architecture, preferential persistence after nuclease challenge, and the immunostimulatory activity of Z-DNA-enriched NET fractions.

## Introduction

DNA is the largest component of neutrophil extracellular traps (NETs) and has traditionally been viewed as a passive structural scaffold, serving to immobilize pathogens and concentrate antimicrobial proteins [1, 2]. This view is increasingly incomplete. NETs occupy a highly dynamic extracellular environment that lacks the regulatory constraints of the intracellular space, exposing their DNA to fluctuating ionic conditions, oxidative stress, and enzymatic remodeling. DNA, in turn, is conformationally plastic, able to adopt noncanonical secondary structures including i-motifs, G-quadruplexes (G4s), and Z-DNA [3]. These structures and their biological roles have been characterized extensively within the nucleus, but little is known about their occurrence in extracellular DNA (eDNA). NET eDNA, moreover, is not an unstructured mass of released chromatin. Atomic force and high-resolution microscopy reveal a branched meshwork of variable pore size and fiber thickness, containing extensively supercoiled naked DNA and a broad heterogeneity in force plateaus indicative of mechanically distinct fiber domains [4]. This architecture is itself proposed to govern the capacity of NETs to physically trap pathogens [4]. NET function may therefore depend not only on how much DNA is released, but on how that DNA is organized once outside the cell. Whether noncanonical conformations are distributed nonrandomly within this network, and whether they relate to its higher-order organization, has not been examined.

Evidence that NET DNA adopts alternative conformations has begun to challenge the view of eDNA as a conformationally uniform scaffold. We recently identified G4 structures within NET eDNA, demonstrating that noncanonical secondary structures can arise within extracellular neutrophil chromatin and carry distinct biochemical properties, including the formation of catalytically active hemin complexes that generate reactive oxygen species and contribute to bacterial killing [5]. In addition, i-motifs were also identified in NETs to selectively bind to thrombin, an interaction that was deemed dependent on both sequence and its noncanonical secondary structure [6]. These observations establish that NET eDNA can contain structurally distinct DNA conformations and raise the possibility that additional noncanonical structures may be intrinsic components of the NET scaffold. Backbone geometry can influence interactions within DNA networks as well as nuclease recognition, and nucleosome loss during NET maturation exposes extended stretches of DNA whose intrinsic conformational properties may become increasingly important [4, 7–9]. DNA conformation can therefore confer properties on NETs beyond those explained by the amount of eDNA alone, and different noncanonical conformations may have distinct structural consequences within the same extracellular chromatin network.

Among noncanonical conformations, Z-DNA has the strongest structural rationale. It is a left-handed helix that exists in dynamic equilibrium with canonical B-DNA [10–12] and is stabilized by torsional stress, high cationic strength, polyamines, and chemical modification [13, 14] — conditions characteristic of inflammatory extracellular environments. Oxidative DNA modifications may further influence this equilibrium, as guanine oxidation to 8-oxoguanine (8-oxoG) alters base-stacking interactions and local DNA flexibility [15]. A structural role for Z-DNA in eDNA networks has precedent in bacteria: during biofilm maturation, eDNA undergoes substantial remodeling and accumulates in the Z-conformation, where it is stabilized by DNABII family proteins and contributes to matrix integrity and resistance to nuclease degradation [16, 17]. Similar principles may apply to NETs, in which DNA is extensively decondensed, exposed to high local concentrations of proteins and cations, and organized into a complex extracellular meshwork. Whether host cells intrinsically generate extracellular Z-DNA, independent of bacterial remodeling, has not been directly examined.

If NETs do contain Z-DNA, the consequences would extend beyond architecture. Mitochondrial DNA (mtDNA) released during NET formation is circular, largely free of nucleosomal packaging, and CpG-rich, representing an additional plausible source of extracellular NET Z-DNA [18, 19]. Oxidative stress has been shown to promote a B-to-Z transition of intracellular mtDNA, enabling sensing by Z-DNA-binding protein 1 (ZBP1) and engagement of receptor-interacting protein kinase 1 (RIPK1) to induce pro-inflammatory molecules in Alzheimer’s disease microglia [20]. This raises the possibility that extracellular Z-DNA within NETs could similarly influence innate immune signaling through ZBP1 or related pathways. Z-DNA is also antigenic, and conformation-specific antibodies have been detected in patients with systemic lupus erythematosus and, at lower levels, in healthy individuals [12]; although microbial eDNA has been proposed as the principal antigenic source, the endogenous contributions sustaining these responses remain undefined.

Here, using conformation-specific immunofluorescence microscopy, quantitative image analysis, and circular dichroism spectroscopy, we identify Z-DNA as an intrinsic structural component of NET eDNA, define its temporal and compartment-specific dynamics during NET progression and maturation, and demonstrate its preferential retention following nuclease treatment. We further examine the relationship between Z-DNA and the higher-order organization of the NET scaffold, quantifying network architecture and topology during NET maturation and following chemical perturbation of the Z:non-Z equilibrium. Pharmacological modulation with chloroquine (CQ) shifts NET DNA toward a non-Z state, remodels network architecture and topology, and reduces the preferential retention of Z-DNA after nuclease challenge. We further find that NET Z-DNA associates with ZBP1 and that Z-DNA-enriched NET fractions elicit greater interferon responses. Together, these findings identify Z-DNA as a dynamic component of NETs across multiple NETosis pathways and associate DNA conformational state with the organization, persistence, and immune activity of extracellular DNA.

## Materials and Methods

### Isolation of human blood neutrophils

Human neutrophils were isolated from peripheral blood obtained from healthy donors following informed consent under a protocol approved by the Stanford University Institutional Review Board (protocol #70759). Neutrophils were isolated using the EasySep™ Direct Human Neutrophil Isolation Kit (#19666, StemCell Technologies, Vancouver, BC, Canada), achieving 99% purity. Following isolation, cells were resuspended in phenol red–free RPMI 1640 medium (#11835030, Gibco, Thermo Fisher Scientific) and used for subsequent experiments.

### *In vitro* neutrophil NETosis stimulation

Freshly isolated neutrophils were seeded in 8-well multi-chamber plates (#PEZGS0816, Millipore) at 1×10 cells/well and allowed to settle for 30 min at 37 °C. NETosis was induced by stimulation with phorbol 12-myristate 13-acetate (PMA, 1 µM; #400145, Cayman Chemical), lipopolysaccharide (LPS, 100 µg/mL; #L2630, Sigma-Aldrich), interleukin-8 (IL-8, 150 ng/mL; #208-IL-050, R&D Systems), tumor necrosis factor-α (TNF-α, 20 ng/mL; #210-TA-20, R&D Systems), or hydrogen peroxide (H□O□, 0.03%; #107209, EMD Millipore) for 4 h, or with the calcium ionophore A23187 (2 µM, #400085, Cayman Chemical) for 1 h, in phenol red-free RPMI-1640 medium (#11835030, Gibco, Thermo Fisher Scientific). Cells were incubated at 37°C, 5% CO□. Unstimulated cells were used as control.

### Preparation of autologous serum and stimulation conditions

Peripheral blood was collected from healthy donors into serum separation tubes (SST). Samples were allowed to clot at room temperature for 20 min and subsequently centrifuged at 1500 × g for 15 min to isolate serum. The resulting autologous serum was collected and used immediately. For serum-based experiments, freshly isolated neutrophils, from the same individual, were resuspended directly in autologous serum without the addition of culture media and stimulated under the indicated conditions as described above.

### Immunofluorescence staining for NET visualization

NETs generated as described above were gently washed once with cold DPBS (#14190250, Gibco, Thermo Fisher Scientific) and fixed with 4% paraformaldehyde (PFA) (#30525-89-4, Thermo Scientific Chemicals) for 20 min at 4°C. Samples were washed three times with DPBS and blocked with eBioscience™ IHC/ICC Blocking Buffer (#00-4953-54, Invitrogen) for 30 min at 37°C.

Immunofluorescence staining was performed using primary antibodies against Z-DNA/Z-RNA (clone Z22) (#Ab00783-23.0, Absolute Antibody), 8-oxoguanine (8-Oxo-G) (#MAB3560, Sigma-Aldrich), and Z-DNA binding protein 1 (ZBP1) (#H00081030-M01, Abnova). To validate PMA-induced NET formation, neutrophils were co-stained with anti-myeloperoxidase (#ab208670, Abcam) and anti-neutrophil elastase (#ab183342, Abcam). In separate MPO and NE co-staining experiments, anti-myeloperoxidase (#MA1-80878, Invitrogen) was used in combination with anti-neutrophil elastase (#ab183342, Abcam). Species-specific fluorophore-conjugated secondary antibodies were selected based on the host species of each primary antibody, including Goat anti-Rabbit IgG (H+L) Alexa Fluor™ 488 (#A-11008, Invitrogen), Goat anti-Mouse IgG (H+L) Alexa Fluor™ 594 (#A-11005, Invitrogen), Goat anti-Rabbit IgG (H+L) Alexa Fluor™ 594 (#A-11012, Invitrogen), Goat anti-Rabbit IgG (H+L) Alexa Fluor™ 647 (#A-21245, Invitrogen) and FITC Donkey anti-Rabbit IgG (minimal x-reactivity) (#406403, BioLegend). Primary antibodies were incubated overnight at 4°C, followed by incubation with secondary antibodies for 1 h at 37°C. To confirm the specificity of Z22 staining, an isotype-matched IgG control (#31235, Thermo Fisher Scientific) was applied at the same final concentration as the Z22 primary antibody and incubated under identical staining conditions. In parallel, secondary antibody-only controls, in which primary antibodies were omitted, were included to assess non-specific secondary antibody binding and background fluorescence.

eDNA was stained with SYTOX (#S11381, Invitrogen) or DAPI (#H-1200-10, VectorLabs). Mitochondrial compartment was labeled using MitoTracker Orange and MitoTracker Green (#M7510 and #M7514, Invitrogen) according to the manufacturer’s instructions over mitochondrial DNA specific dyes to avoid any confounding effect on B/Z states. Fluorescence images were acquired using a Nikon ECLIPSE Ti2-E microscope and analyzed either using ImageJ software or custom python pipeline described below.

### NET DNA preparation

NETs were generated as described above. To isolate NETs DNA for spectroscopic analyses, samples were treated with AluI restriction enzyme (#ER0011, Thermo Fisher Scientific) at 10 U per well for 30 min at 37 °C. Following digestion, the supernatant from each well was collected and centrifuged at 300 × g for 5 min at 4 °C to remove intact cells and cellular debris. The resulting supernatant containing NET-derived DNA was used for downstream structural analyses.

### Biophysical characterization of NET-derived DNA

#### a. Circular dichroism (CD) spectroscopy

NETs DNA samples were analyzed using CD spectroscopy to evaluate DNA conformational signatures. CD spectra were recorded using a quartz cuvette with a 1 mm path length over the wavelength range of 250–300 nm, and ellipticity values were collected in millidegrees (mdeg). The scanning time was set to 0.5 s per point, and each spectrum represents the average of three scans. DPBS buffer was used as the blank control.

As structural controls, poly(dG–dC) duplex DNA (#P9389, Sigma-Aldrich) was analyzed in both B-DNA and Z-DNA conformations. The Z-DNA conformation was induced by incubating poly(dG–dC) in 4 M NaCl (#AM9759, Invitrogen) for 1 h at room temperature, whereas untreated poly(dG–dC) served as the B-DNA control. These controls were analyzed under identical CD conditions to validate the spectral signatures observed in NET-derived DNA samples. The obtained spectra were compared and further analyzed using Chirakit, the online tool for analysis of CD spectroscopy data[21]. Briefly, given that the NET-derived samples were a mixed population containing proteins, antimicrobial peptides, DNA, polyamines, and other molecules, and that our focus was solely on the DNA, we narrowed the wavelength to between 250 to 300 nm. Next, we performed spectra decomposition using singular value decomposition and principal component analysis to reduce the DNA constituents of the NET population into a set of basis spectra. The derived basis spectra were then compared against the reference spectra for canonical B and non-canonical Z DNA after L2-normalization to generate spectral comparison plots and estimate Euclidean distances to assess similarity to the reference spectrum [22, 23].

#### b. UV absorbance analysis

DNA absorbance spectra were measured using a NanoDrop spectrophotometer. The A295/A260 ratio was used as an indicator of DNA conformation [24]. Ratios were compared against poly(dG-dC) controls prepared in B- and Z-DNA conformations to further validate the presence of Z-DNA conformations.

#### c. Assessment of nuclease sensitivity

NETs DNA samples were subjected to digestion with DNase I (#EN0521, Thermo Fisher Scientific), DNase1L3 (#HY-P71543, MedChemExpress), RNase A (#EN0531, Thermo Fisher Scientific), or S1 nuclease (#EN0321, Thermo Fisher Scientific) (1U each, 37°C, 15 min).

The nuclease reactions were terminated by the addition of EDTA (#R1021, Thermo Fisher Scientific) to a final concentration of 10 mM, followed by heat inactivation at 65°C for 10 min using a heat block.

### Modulation of oxidative stress using vitamin C (L-ascorbic acid)

Neutrophils were pre-treated with vitamin C (L-ascorbic acid, #A0278, Sigma) at a final concentration of 2 mM for 30 min prior to stimulation and maintained throughout NETosis.

### Cationic modulation of DNA conformation

To promote conditions favoring the B-to-Z DNA conformational transition, NETs formed following 3 h of PMA (1 µM) stimulation of neutrophils from healthy donors were treated with spermidine trihydrochloride (#S2501, Sigma-Aldrich) at final concentrations of 0.5, 1, 2, 4, or 8 mM or or CaCl (#C1016, Sigma-Aldrich) at final concentrations of 1, 2, 4, or 8 mM, during the final 1 h of the total stimulation period. This timing ensured that spermidine and CaCl were introduced after the initiation of NETosis. Following treatment, NETs were subjected to fluorescence imaging as described above.

### Pharmacological modulation of DNA conformation

To induce Z-to-B DNA conformational transition, NETs formed following 3 h of PMA stimulation of neutrophils from healthy donors were treated with chloroquine (10 µM, #C6628, Sigma-Aldrich) during the final 1 h of the total stimulation period. This timing ensures that chloroquine was introduced well after the initiation of NETosis. Following treatment, NETs were subjected to nuclease digestion assays and structural analysis as described above.

### Immunostimulatory characterization of NET-derived DNA

#### A. Immunoprecipitation of Z-DNA-enriched NET fractions

NET-derived DNA was prepared from PMA-stimulated neutrophils following AluI digestion, as described above. Clarified NET DNA-containing supernatants were subjected to immunoprecipitation using the Z-DNA/Z-RNA-specific Z22 antibody (#Ab00783-23.0, Absolute Antibody) and the Pierce™ MS-Compatible Magnetic IP Kit (Protein A/G; #90409, Thermo Fisher Scientific), following the manufacturer’s protocol with a minor modification at the elution step. Briefly, magnetic Protein A/G beads were equilibrated, coupled to Z22 antibody, and incubated with NET-derived DNA samples to capture Z22-reactive material. Following magnetic separation, the unbound supernatant was retained as the non-Z-DNA fraction. Beads were washed according to the kit protocol, and Z22-bound material was eluted using DNA elution buffer consisting of 10 mM Tris-HCl pH 8.0 and 0.1 mM EDTA, in place of the kit-provided elution buffer. DNA concentrations in the Z-DNA-enriched and non-Z-DNA fractions were quantified using a NanoDrop spectrophotometer and normalized prior to downstream functional assays.

#### B. PBMC isolation and stimulation with NET-derived DNA fractions

Human peripheral blood mononuclear cells (PBMCs) were isolated from the same three healthy donors peripheral blood obtained at multiple independent timepoints (n=11 paired observations) using the EasySep™ Direct Human PBMC Isolation Kit (#19654, STEMCELL Technologies), according to the manufacturer’s instructions. Isolated PBMCs were washed and resuspended in phenol red–free RPMI 1640 medium (#11835030, Gibco, Thermo Fisher Scientific) and seeded in 12-well culture plates at 1 × 10 cells per well. Cells were stimulated with autologous Z-DNA-enriched or corresponding non-Z-DNA NET-derived DNA fractions at final DNA input amounts of 2 µg or 5 µg per condition. Lipopolysaccharide (LPS, 100 ng/mL; #L2630, Sigma-Aldrich) was included as an inflammatory stimulation control, and unstimulated cells were included as the cells-only background control. Cells were incubated for 24 h at 37°C and 5% CO□. Following stimulation, culture supernatants were collected and centrifuged to remove residual cells and debris before IFN-α quantification.

#### C. IFN-α ELISA

IFN-α concentrations in 11 paired PBMC culture supernatants were quantified using the Human IFN alpha ELISA Kit (#BMS216, Invitrogen, Thermo Fisher Scientific), according to the manufacturer’s instructions. These observations reflected measurements across three independent donors at multiple timepoints and doses, capturing both inter-and intra-donor variability. Samples and standards were added to the ELISA plate, followed by sequential incubation withdetection reagents and substrate solution as specified by the kit protocol. The reaction was stopped using the kit-provided stop solution, and absorbance was measured at 450 nm using a microplate reader. IFN-α concentrations were calculated from the standard curve and background-normalized to cells-only controls. For fold-change analyses, IFN-α responses induced by Z-DNA-enriched fractions were compared with their corresponding non-Z-DNA fractions at matched DNA input amounts.

### Image Processing and Quantitative Analysis of NET DNA Structure

Fluorescence microscopy images were analyzed using a custom Python-based pipeline to quantify the structural organization and composition of eDNA during NETosis and NET maturation. The pipeline was built on various libraries including scikit-image, SciPy, numpy, matplotlib, tifffile, and pandas [25–28]. Multichannel images were separated into individual non-Z-DNA and Z-DNA channels for independent processing. All analyses were performed on raw images without prior thresholding or manual preprocessing. The workflow is represented in Supplementary Fig. S1.

#### A. Preprocessing and Segmentation

Each channel was first normalized to its dynamic range and background-corrected using percentile-based scaling. To identify eDNA structures, a fixed intensity thresholding was applied following normalization in combination with edge-based (Sobel) filtering to suppress diffuse background signal and enhance filamentous features. Binary masks were further refined using morphological and size filtering to remove small, isolated objects and retain filamentous structures. Images generated across all experimental conditions and channels were processed using identical processing logic and parameter settings to ensure consistency. Given the lower signal-to-noise ratio of Z-DNA staining, segmentation parameters were tuned to prioritize high-confidence, filament-associated signal while minimizing inclusion of background noise, thereby resulting in structurally resolved DNA rather than total fluorescence intensity. Filamentous DNA structures were defined based on morphological criteria including elongation and minimum size constraints. Quantitative fluorescence microscopy coupled with computational image analysis has been established for objective assessment of NET formation and morphology, providing alternatives to subjective manual quantification[29, 30]. We therefore applied a standardized multichannel image-analysis pipeline to resolve DNA architecture and quantify the spatial organization and structural contribution of Z-DNA within the NET network.

#### B. Skeletonization and Structural Representation

Binary masks were skeletonized to obtain one-pixel-wide representations of DNA filaments. This transformation preserved the topology and connectivity of extracellular DNA while enabling quantitative analyses of filament length and organization independent of thickness or intensity. Skeleton length can describe the NETs filamentous architecture and geometry more appropriately than pixel intensity which can be confounded by local antibody access, dye penetration, and imaging depth. [4, 31–33] NET geometry and topology were quantified from the segmented masks and skeletons, including network area and length, filament density and thickness, branch points, endpoints, junction density, triple- and quadruple-point frequencies, fractal dimension, lacunarity, and pore number and geometry. These metrics were used to characterize higher-order NET architecture and to assess its relationship with Z-DNA abundance and distribution during NET maturation and following chemical perturbation of DNA conformation. Pores were identified as enclosed regions within the segmented NET mask and quantified by area, equivalent diameter, aspect ratio, circularity, and solidity. For temporal analyses, the relationships between Z-DNA and NET topology were further assessed using standardized multiple linear regression of Z-DNA and residual non-Z-DNA skeleton content against independent measures of network complexity.

#### C. Quantification of Z-DNA Organization and Nucleation

Z-DNA organization was quantified using two complementary metrics: (i) the fraction of Z-DNA that is spatially independent of non-Z-DNA signal, and (ii) the number of discrete Z-DNA domains. Z-DNA domains were defined as spatially contiguous regions within the Z-DNA mask, identified using connected-component labeling, a standard image-analysis method that groups adjacent foreground pixels into discrete objects, aligned on the non-Z-DNA mask. Domain counts were used as a proxy for the appearance of spatially discrete Z-DNA structures; this image-based metric does not directly resolve molecular nucleation events.

#### D. Intensity and Enrichment Measurements

Total and mean fluorescence intensities were computed for each channel within the segmented extracellular DNA regions. Z-DNA enrichment relative to non-Z-DNA was quantified using the logarithm of the ratio of Z-DNA to non-Z-DNA integrated intensity. Intensity measurements were normalized to the NET area where appropriate to account for differences in DNA spread and density.

#### E. Spatial and Functional Association Analyses

Colocalization and spatial proximity between Z-DNA and other molecular markers (including ZBP1, MitoTracker, and 8-oxoG) were quantified using overlap-based metrics (Manders’ coefficients and Jaccard index) and distance-based measures. These analyses were performed on segmented masks to ensure that comparisons reflected structured signal rather than diffuse background. Z-DNA association with pore boundaries was additionally assessed by quantifying the fraction of pore-associated boundary pixels containing Z-DNA and the proportion of total pore area associated with Z-DNA-positive boundaries. Z-DNA spatial distribution relative to NET topology was assessed by classifying Z-DNA skeleton and intensity signal according to proximity to B-DNA skeleton junction and endpoint nodes (junction-proximal, endpoint-proximal, non-node) and by local network thickness (thin, intermediate, thick tertiles), using the non-Z-DNA (B-DNA) skeleton as the structural reference. To assess the sensitivity of this classification to segmentation parameters, the analysis was repeated with a fixed Sobel edge-detection threshold calibrated from the pooled gradient distribution across the full image set, in place of the threshold recomputed independently for each image in the primary analysis; the B-DNA channel was analyzed in parallel as a specificity control. For ZBP1 and 8-oxoG analyses, partial correlation coefficients controlling for local SYTOX intensity were additionally computed to account for differences in local DNA abundance. The spatial relationship between 8-oxoG and Z-DNA was further evaluated as a function of distance from Z-DNA-positive regions. NET-constrained randomization (permutation of ZBP1 signal within the segmented NET skeleton) was used to test whether observed ZBP1–Z-DNA spatial association exceeded that expected from NET geometry and signal abundance alone. ZBP1 analyses were restricted to images passing predefined quality-control criteria excluding images with a Z-DNA-positive skeleton compartment below a minimum size threshold, and mitochondria-stratified comparisons were further restricted to images with an available MitoTracker channel and sufficient mitochondrial and non-mitochondrial Z-DNA-positive pixels for comparison.

#### F. NET Progression State Definition

To capture the progression of NET formation, individual samples were assigned to discrete NET states (Early, Mid, Late) based on a composite progression score. This score combined normalized stimulation timepoint (empirically weighted 0.65) with non-Z-DNA (SYTOX) skeleton length normalized within each timepoint (empirically weighted 0.35), and samples were divided into tertiles of this composite score to assign Early, Mid, and Late NET categories, enabling classification of samples along a continuum of NET maturation, independent of Z-DNA structural metrics. To assess robustness of the NET state classification, the analysis was repeated using timepoint alone as the progression score (α=1.0). Classification agreement with the composite score was estimated and Kruskal-Wallis test was performed to compare the medians between the two groups for each state.

#### G. Statistical Analysis

All statistical analyses were performed using non-parametric methods unless otherwise specified. Paired comparisons were conducted using Wilcoxon signed-rank tests or paired t-tests, and group comparisons were performed using Mann–Whitney U tests. Correlations were assessed using Spearman’s rank correlation coefficient. Effect sizes were calculated where appropriate. All reported trends were supported by multiple independent metrics to ensure robustness against segmentation or measurement bias.

## Results

### Z-DNA is an intrinsic structural component of eDNA in NETs

To determine whether Z-DNA constitutes an intrinsic component of NET-derived eDNA, we performed immunofluorescence staining using a Z-DNA–specific antibody (clone Z22) in neutrophils under basal and NETosis-inducing conditions, namely 1 µM PMA stimulation for 4 hours. Recent structural studies have demonstrated that Z22 specifically recognizes the characteristic left-handed backbone geometry of Z-DNA in a sequence independent manner, while multiple independent studies have successfully combined Z22 with orthogonal approaches, including ChIP-seq, biochemical assays, and genetic perturbations, to investigate endogenous Z-DNA without cross reactivity [8, 34–36]. NET formation was visualized using SYTOX Red, a nucleic acid-binding dye that produces little or no fluorescence when interacting with Z-DNA; SYTOX Red signal in this context therefore reflects non-Z-DNA regions of the NET scaffold [37] (Fig. 1A). In unstimulated neutrophils, Z-DNA signal was minimal and largely confined to intact nuclei, with no detectable eDNA. In contrast, PMA stimulation induced robust NET formation, confirmed by co-staining with myeloperoxidase and neutrophil elastase decorating the NETs (Supplementary Fig. S2), characterized by extensive SYTOX Red–positive filamentous eDNA structures accompanied by a pronounced increase in Z-DNA signal that closely traced the eDNA network, accounting for approximately 20% of the integrated eDNA fluorescence signal (Fig. 1A). This was morphologically distinct from the nuclear condensation and fragmentation observed with apoptosis, and necrosis, in which loss of membrane integrity occurs while the nucleus initially remains relatively intact [38, 39]. Isotype and secondary antibody only controls confirmed the specificity of the Z22 antibody and verified true detection of Z-DNA in NETs (Supplementary Fig. S3). Notably, Z-DNA signal followed elongated, web-like fibers, indicating its direct incorporation into NET-derived eDNA. Additionally, to assess whether Z-DNA formation is dependent on a specific NETosis pathway, we next examined stimuli that engage distinct mechanisms, including another classical suicidal NETosis inducer (H□O□), inflammatory mediators (TNF-α, IL-8, and LPS), and A23187, a calcium ionophore that triggers rapid, NOX-independent, mtDNA–rich vital NETosis [40–42]. Despite these mechanistic differences, all conditions consistently exhibited Z-DNA formation in the eDNA, although skeletonization based image analysis showed variability in the total Z-DNA length based on the stimulus (Fig. 1B and Supplementary Fig. S4). To assess Z-DNA formation under physiologically relevant conditions, we performed an *ex vivo* assay where neutrophils were stimulated in the presence of autologous serum. Z-DNA was observed in the NETs across the different stimuli *ex vivo* as well but due to nutrient limitation there was an overall drop in the number of viable neutrophils and therefore the NETs signal (Supplementary Fig. S5). In addition, consistent with the established Z-DNA-stabilizing properties of polyamines, titrated addition of spermidine was associated with an overall positive trend in Z-DNA enrichment across the tested concentration range (Z/non-Z-DNA skeleton ratio: ρ = 0.49, p = 0.052; Z/non-Z-DNA intensity ratio: ρ=0.918, p=5.50×10□□). Increasing spermidine concentration was also associated with substantial remodeling of NET architecture, characterized by increased NET thickness (ρ = 0.68, p = 0.0037), together with decreased junction density (ρ = −0.55, p = 0.027), triple-point frequency (ρ = −0.56, p = 0.024), quadruple-point frequency (ρ = −0.59, p = 0.016), and filament density (ρ = −0.52, p = 0.038), while NET area remained unchanged (ρ = 0.08, p = 0.77; Supplementary Fig. S6). We also used CaCl as a chemically distinct perturbation of DNA conformation. Similar to spermidine, increasing CaCl concentration produced a strong concentration-associated increase in Z-DNA enrichment (Z/non-Z-DNA skeleton ratio: ρ = 0.80, p = 0.0018; Z/non-Z-DNA intensity ratio: ρ=0.928, p=1.31×10□^5^) and was accompanied by increased NET thickness (ρ = 0.58, p = 0.047). In contrast, NET area showed no concentration-dependent change (ρ = 0.13, p = 0.69), while junction density (ρ = −0.41, p = 0.19), triple-point frequency (ρ = −0.45, p = 0.14), quadruple-point frequency (ρ = −0.37, p = 0.24), and filament density (ρ = −0.22, p = 0.50) showed concordant but non-significant downward trends (Supplementary Fig. S7). Both chemically distinct perturbations resulted in convergent architectural consolidation of the NET meshwork resulting from a combination of increased thickness and reduced junctional and branching metrics without affecting the overall area. Our findings across the most commonly used *in vitro* stimulants of vital and suicidal NETosis, and sterile inflammations indicate that Z-DNA generation represents a convergent feature of NETosis across distinct pathways, agnostic of stimulus suggesting an integral structural role of the non-canonical form of DNA.

**Figure 1:**
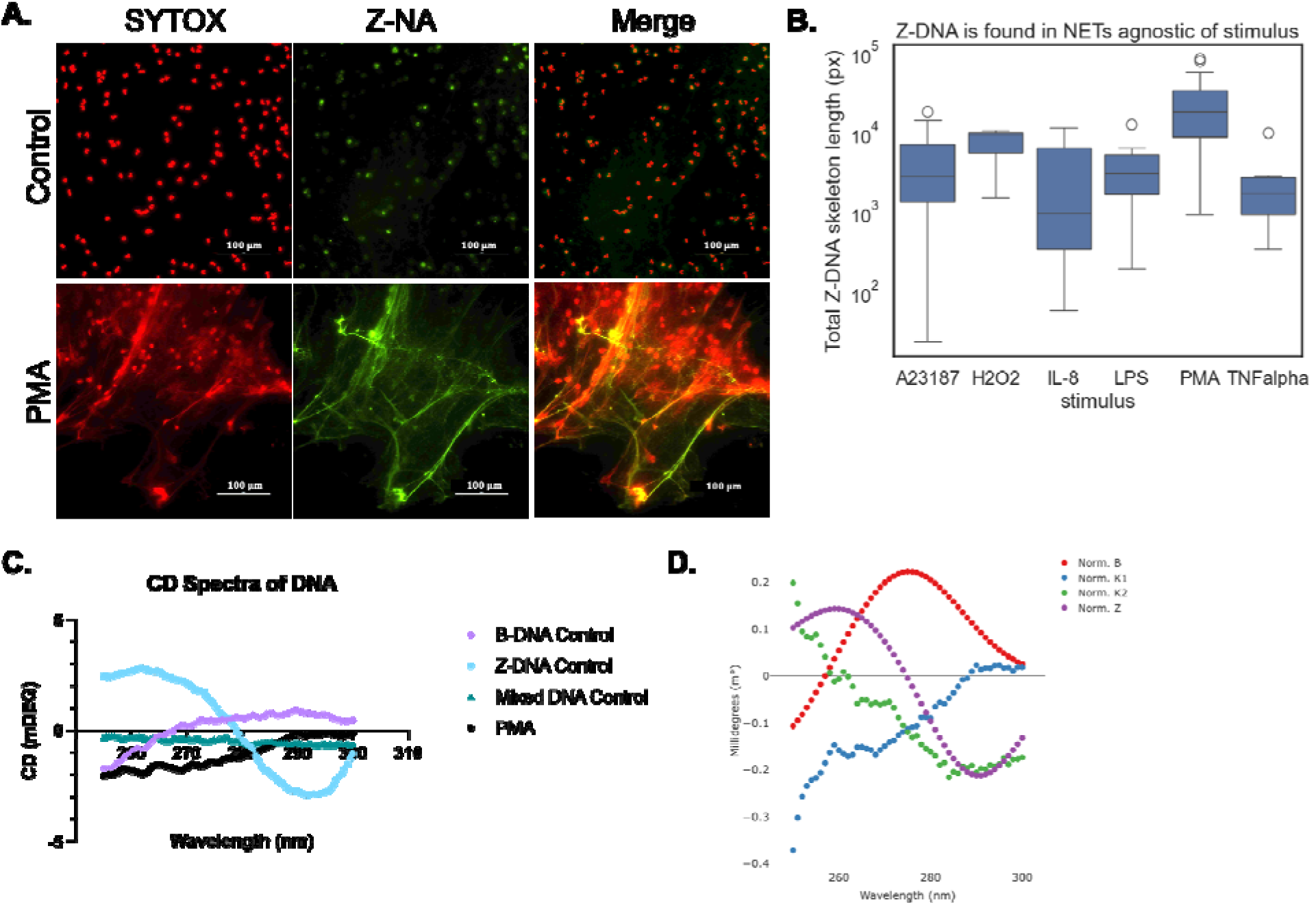
Z-DNA is intrinsic to NETs eDNA. **(A)** Representative immunofluorescence image of neutrophils under unstimulated conditions and following PMA-induced (1 µM, 4h) NETosis. Z-DNA was detected using a Z-DNA–specific antibody (clone Z22) (green) and non-Z-DNA wa stained with SYTOX (red). Merged images show co-localization of Z-DNA with total DNA structures. Scale bars, 100 µm **(B)** Quantitative analysis of total Z-DNA skeleton length measured from 102 images across 6 different stimulation conditions, including PMA, LPS, A23187, TNF-α, IL-8, and H O showing widespread Z-DNA presence irrespective of inducing stimulus. **(C)** Orthogonal, antibody free confirmation of Z-DNA presence in NETs. CD spectra of NET-derived eDNA following PMA-induced NETosis, shown alongside reference spectra of poly(dG–dC) in canonical B-form, salt-induced Z-form, and mixed conformations. **(D)** Spectral comparison of deconvoluted NETs basis spectra against reference DNA structural signatures, indicating the degree of resemblance of NET-derived eDNA to B-DNA and Z-DNA, confirming the presence of Z-DNA in the mixed populations of NETs eDNA.

To further confirm the existence of Z-DNA specifically in NETs, we treated PMA NETs with RNase A and S1 nuclease. Treatment with RNase A reduced total extracellular signal proportionally across both SYTOX and Z22 channels (Z/non-Z-DNA intensity ratio 0.31 vs. 0.35; Z/non-Z-DNA skeleton ratio 0.55 vs. 0.48), indicating that RNase does not selectively deplete Z22 signal and that RNA is not a major contributor to the Z22-positive structures. This interpretation is consistent with prior evidence that Z-RNA is susceptible to RNase A degradation, which would be expected to eliminate the Z22 signal if the conformation were RNA-based [43]. In contrast, S1 nuclease treatment markedly reduced eDNA signal, with total non-Z-DNA signal reduced by 49% and Z-DNA signal reduced by 72% (Supplementary Fig. S8). This disproportionate reduction of the Z22 signal indicates that Z-DNA–positive NET regions are preferentially sensitive to S1 nuclease compared with bulk non-Z-DNA. This interpretation is consistent with recent work showing that while DNase I lacks activity against Z-DNA, S1 nuclease can degrade Z-DNA in eDNA matrices [8], further supporting the conclusion that these Z22-positive structures abundant in NETs released in response to various stimuli indeed represent Z-DNA.

To orthogonally validate the existence of Z-DNA within the conformational ensemble of NETs eDNA in an antibody independent manner, we performed circular dichroism (CD) spectroscopy on NET-derived eDNA following PMA induction (1 µM, 4 h). Spectral analysis revealed features consistent with non-B DNA conformations, including a partial negative ellipticity near ∼295 nm, a characteristic signature of Z-DNA [44]. This spectral feature is distinct from other noncanonical nucleic acid structures previously described in NETs, including G4, which characteristically produces positive ellipticity in this region regardless of topology, and i-motifs, which exhibit a characteristic positive band near 285–290 nm and lack the inverted Z-form signature [45, 46].These profiles were supported by poly(dG–dC) controls in canonical B-form, salt-induced Z-form and mixed conformations, confirming the presence of Z-DNA within a heterogeneous eDNA population (Fig. 1C). Consistent with this, singular value decomposition of the NETs eDNA CD spectra into basis spectra and subsequent spectral comparison against reference signatures of canonical B-DNA and Z-DNA revealed that basis spectrum k2 aligned with Z-DNA (Fig. 1D) with a Euclidean distance of 0.8. Interestingly, basis spectrum k1 did not align perfectly with the reference spectrum for B-DNA with a Euclidean distance of 1.8, requiring further resolution and reinforcing the notion that NETs eDNA exists in structurally diverse states. Although NET-derived eDNA represents a complex mixture containing associated proteins and polyamines, protein secondary structure signals are predominantly confined to the far-UV region below 250 nm, whereas CD features arising from DNA conformational transitions are characteristically observed above 250 nm, supporting that the spectral signatures detected within the 250–300 nm window analyzed here primarily reflect DNA backbone conformation rather than protein structural contributions [47, 48].

### Temporal and compartment-specific dynamics of Z-DNA formation during NETosis

We next sought to investigate the temporal dynamics of Z-DNA formation and integration in NETs during NETosis. Time-resolved imaging every hour, over 6 hours, following PMA stimulation revealed a staged evolution and progression in the source and distribution of Z-DNA in the extracellular space (Fig. 2A). At early time points, eDNA release was limited; however, Z-DNA signal was already detectable in regions aligned with mitochondrial compartment labeling, stained with MitoTracker, indicating early mt-associated Z-DNA release. As NETosis progressed, extensive filamentous DNA structures emerged, accompanied by a marked increase in Z-DNA signal along these networks. At later stages, Z-DNA became predominantly associated with non-mt-associated DNA, nuclear eDNA, while mt-associated Z-DNA remained detectable, indicating a transition from early mitochondrial release to broader nuclear DNA contribution. This is readily apparent in the quantitatively estimated total Z-DNA length (in pixels) over time in the mt-associated Z-DNA and non-mt-associated Z-DNA measured across multiple replicate images over time (Fig. 2B). Mt-associated Z-DNA–associated skeleton length remained relatively stable across time, whereas the non-mt-associated Z-DNA component increased substantially, resulting in a progressive divergence between compartments. These data indicate that the expansion of Z-DNA during NETosis is dominated by increasing nuclear DNA–associated structures, superimposed on an early, persistent mt-associated DNA-derived component. Of note, to our knowledge this is the first demonstration of extracellular mt-associated DNA signals, non-Z- or Z-DNA, released early during PMA-stimulated NETosis, a feature classically associated with vital NETosis pathways as attributed to A23187 stimulation (Supplementary Fig. S9B). This finding suggests that early mt-associated DNA release may present a secondary point of convergence between mechanistically distinct NETosis pathways. Complete time-resolved datasets across all time points and stimuli are provided in Supplementary Fig. S9.

**Figure 2:**
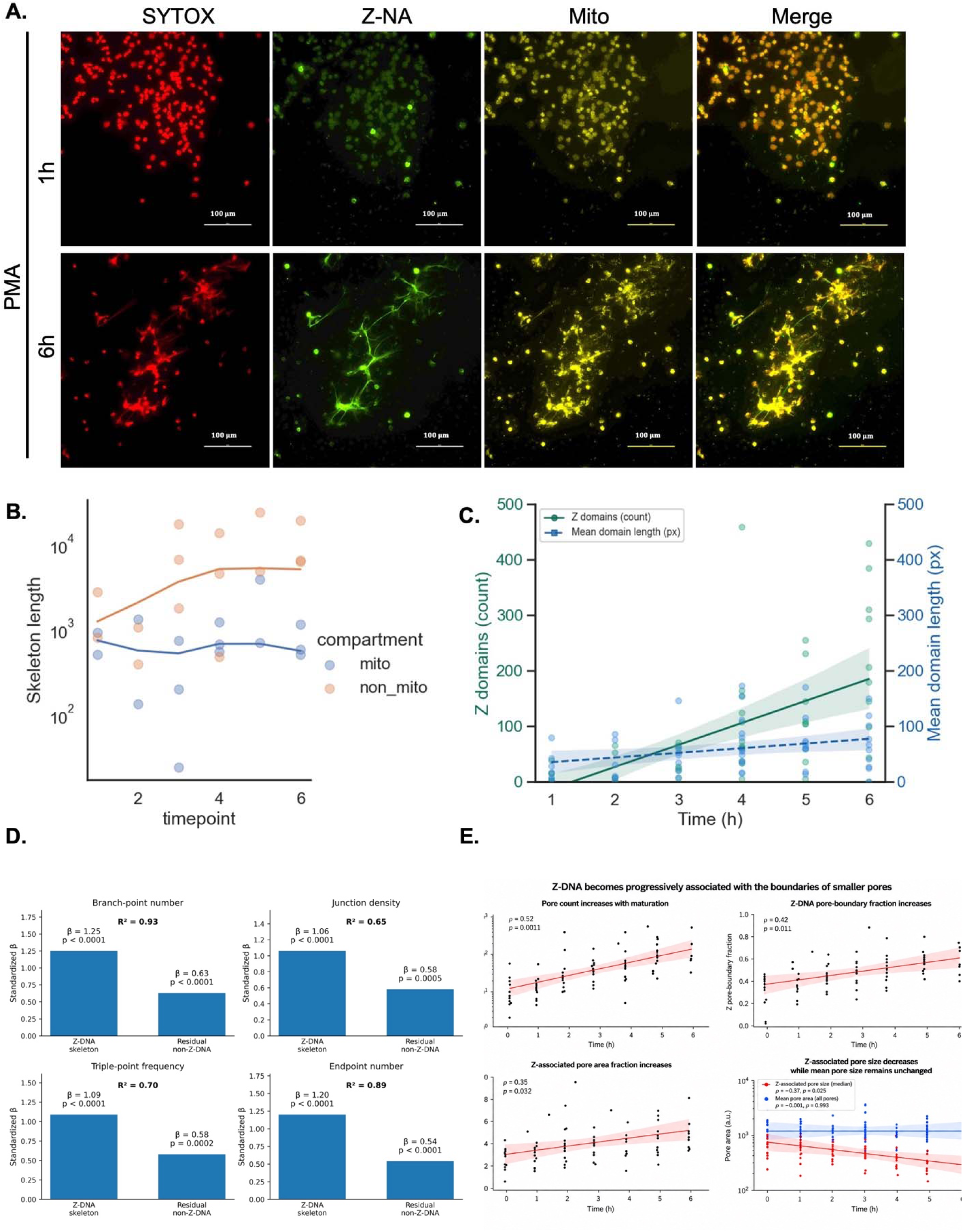
Time-course analysis of Z-DNA formation dynamics and its disproportionate association with topological remodeling as NETs mature. **(A)** Representative immunofluorescence images of neutrophils undergoing PMA induced NETosis (1h) and late time points (6h), stained for Z-DNA (Z-NA, green), mitochondrial compartment (MitoTracker, yellow), and non-Z-DNA (SYTOX, red), with corresponding merged images Scale bars, 100 µm. **(B)** Quantification of total Z-DNA skeleton length over time during PMA induced NETosis measured from ∼8 images/time point. Total Z-DNA skeleton length per field of view is shown, stratified by mt-associated DNA (Mitotracker channel in ∼3 images/time point) and non-mt-associated DNA. Each point represents an independent field of view; lines indicate locally weighted scatterplot smoothing trend over time. Early enrichment of mt-associated Z-DNA which plateaus over time while non-mt-associated DNA steadily increases over time. **(C)** Number of Z domains and the mean domain length over time. Number of Z domains increases over time while the mean domain length stays relatively consistent through time. Shaded regions indicate the 95% confidence interval. **(D)** Standardized regression coefficients comparing the associations of Z-DNA and residual non-Z-DNA skeleton components with branch-point number, junction density, triple-point frequency, and endpoint number. Z-DNA showed consistently stronger associations with each topological feature. **(E)** Relationships between NET maturation and pore architecture. Pore number increased with maturation, accompanied by increases in the Z-DNA-associated pore-boundary and pore-area fractions, while the median size of Z-DNA-associated pores decreased; mean pore area across all pores remained unchanged. Together, these findings indicate that Z-DNA accumulation is preferentially associated with increasing network complexity and remodeling of the pore architecture during NET maturation.

To determine whether the increase in total Z-DNA length over time was associated primarily with the appearance of additional spatially discrete Z-DNA domains rather than elongation of existing domains, we defined Z domains as discrete, spatially contiguous connected components within the segmented Z-DNA mask, representing individual Z-DNA units. We assessed the robustness of our domain detection by estimating the relationship between the domain count and signal intensity. Z-DNA domain count positively correlated with both Z-DNA (r = 0.43, r² = 0.19, p < 10□□) and non-Z-DNA (r = 0.54, r² = 0.29, p < 10□□) signal. While these relationships are modest in strength, their consistency across two independent channels is consistent with domain formation scaling with NET DNA abundance rather than arising as an artifact of low signal fragmentation; the unexplained variance likely reflects additional contributions from domain size and distribution, examined further below. Consistent with the observed progression in the total Z-DNA skeleton length over time, there exists a time-dependent increase in the formation of new Z-DNA domains, as reflected by an increase in Z-DNA domains while the mean domain length plateaus over time (Fig. 2C). In fact, ordinary least squares regression analysis (n = 49 fields; full model R² = 0.82, adj. R² = 0.80, F = 50.19, p = 7.65×10□¹□) determined that the number of Z-DNA domains (β = 62.0, 95% CI [20.0, 104.1], p = 0.005) and its interaction with mean domain length (β = 0.70, 95% CI [0.11, 1.30], p = 0.022) were significant predictors of total Z-DNA structure in NETs, while mean domain length alone was not (β = −11.5, 95% CI [−75.9, 52.9], p = 0.72). Together, these results establish that Z-DNA formation is a dynamic and continuously evolving process, driven by ongoing nucleation and structural expansion rather than passive redistribution and elongation of previously formed Z domains.

### Z-DNA dynamics over NETs progression

To obtain a global depiction of NET DNA architecture progressing over time, encompassing both non-Z and Z-DNA exclusively in the extracellular regions, we first developed a NET state classification system. NET progression states (Early, Mid, Late) were defined using a composite score integrating normalized timepoint and relative non-Z-DNA structural expansion, capturing the continuum of NET maturation (Supplementary Fig. S10A). Such classification further corroborated our previous findings and shows significant increases in Z domains as well as the Z-DNA skeleton length in the progression of NETs (Supplementary Figs. S10B and C). Interestingly, the fraction of Z-DNA independent of the non-Z-DNA backbone is significantly enriched in the early states (Early vs Late; p = 5 x 10□^6^). This trend was not dependent on state binning, as Z-DNA structural independence decreased continuously with normalized time (r = −0.47, p < 10□□), while showing minimal dependence on the structural component used to define the classifier, indicating that observed dynamics reflect true progression rather than binning artifacts. Furthermore, classification agreement between timepoint alone and composite score was 78.2%, and Z-DNA skeleton length and domain count trajectories remained highly significant across all NET progression states (both classifiers p<0.0001), confirming that observed Z-DNA progression is not an artifact of classifier construction. Integrating the data, overall while mt-associated Z-DNA provides an initial efflux of Z-DNA, enriched in the Early NETs, as eDNA becomes more abundant in the extracellular space, environmental factors, NETs associated proteins, polyamines, antimicrobial peptides, or other unknown factors drive nucleation of new Z domains as NETs progress.

Importantly, increasing absolute Z-DNA abundance was not accompanied by a monotonic increase in the overall Z:non-Z-DNA composition of the extracellular scaffold. Across the PMA time series, Z-DNA skeleton length increased progressively with time (ρ = 0.65, p = 2.16 × 10□¹¹), whereas the Z:non-Z-DNA skeleton ratio showed no significant temporal trend (ρ = −0.13, p = 0.26). Thus, Z-DNA accumulation during NET maturation reflects increasing incorporation of Z-DNA into an expanding extracellular DNA network rather than a simple progressive conversion of the NET scaffold toward a Z-DNA-dominant state.

We next asked whether the accumulation of Z-DNA during NET maturation simply tracked expansion of the eDNA network, or whether Z-DNA was preferentially incorporated into specific features of the developing topology. Across the PMA time series, Z-DNA skeleton content showed a consistently stronger association with multiple independent measures of network topology than the residual non-Z-DNA skeleton component. Z-DNA was more strongly associated with branch-point number (β = 1.25, p < 0.0001) than the residual non-Z-DNA component (β = 0.63, p < 0.0001), with the model explaining 93% of the variance (R² = 0.93). Similar relationships were observed for junction density (βZ = 1.06, p < 0.0001; βB = 0.58, p = 0.0005; R² = 0.65), triple-point frequency (βZ = 1.09, p < 0.0001; βB = 0.58, p = 0.0002; R² = 0.70), and endpoint number (βZ = 1.20, p < 0.0001; βB = 0.54, p < 0.0001; R² = 0.89). Consistent with this compositional remodeling, total network length increased with maturation (ρ = 0.77, p < 0.0001), as did branch-point number (ρ = 0.72, p < 0.0001), junction density (ρ = 0.69, p < 0.0001), triple-point frequency (ρ = 0.69, p < 0.0001), and quadruple-point frequency (ρ = 0.69, p < 0.0001), while filament thickness decreased (ρ = −0.76, p < 0.0001) (Fig. 2D) consistent with decondensation of chromatin as NETs mature. Pore number also increased (ρ = 0.51, p = 0.0011) with an increase in Z-DNA associated pore-boundary fraction (ρ = 0.42, p = 0.011) and Z-DNA associated pore area fraction (ρ = 0.35, p = 0.032) but decrease in median Z-DNA associated pore size (ρ = −0.37, p = 0.025) while mean pore area across all pores remained unchanged (Fig. 2E). Together, these findings indicate that Z-DNA is not simply a passive readout of increasing eDNA abundance but is disproportionately associated with the topological features that emerge as the NET network matures. These data are consistent with Z-DNA becoming increasingly integrated into the developing NET scaffold, describing a transition from a relatively compact eDNA structure toward a larger, thinner, and increasingly interconnected meshwork, with more branches, junctions, and discrete pores, while Z-DNA becomes increasingly incorporated into this developing architecture.

### ZBP1 associates with extracellular NETs Z-DNA

Given our discovery of mitochondria-associated Z-DNA in PMA-stimulated NETs, we asked whether these extracellular Z-DNA structures associate with ZBP1, a known intracellular sensor and stabilizer of Z-form nucleic acids. Immunofluorescence analysis revealed redistribution of ZBP1 to eDNA structures following PMA stimulation, where ZBP1 signal was frequently aligned with Z-DNA-positive regions along eDNA fibers (Fig. 3A). This association was also observed in regions enriched for mt-associated Z-DNA. Following A23187 stimulation, ZBP1 similarly colocalized with Z-DNA-positive eDNA structures containing mt-associated signal (Supplementary Fig. S11A). Pixel-based colocalization analysis showed greater overlap of ZBP1 with mt-associated Z-DNA than with non-mt-associated Z-DNA, including a higher fraction of mt-associated Z-DNA overlapping ZBP1 (Manders’ M2: 0.39 vs. 0.04; Wilcoxon p = 5.3 × 10□□). ZBP1 also showed greater overlap with Z-DNA than with non-Z-DNA (p = 0.01). Consistent with these observations, analysis of SYTOX-defined NET skeletons showed higher ZBP1 signal within Z-DNA-positive regions relative to Z-DNA-negative regions (Fig. 3B). Across quality-controlled images, this enrichment was significant in both PMA-induced NETs (median 1.5x, range 0.61–2.17x, n = 17; Wilcoxon p = 0.0056) and A23187-induced NETs (median 1.23x, range 0.20–2.02x, n = 17; Wilcoxon p = 0.0093, Supplementary Fig. S11B). However, NET-constrained randomization showed that the magnitude of this association was not significantly greater than that expected from NET geometry alone when assessed across the image population (PMA median enrichment 1.03-fold, p = 0.225; A23187 median 1.08-fold, p = 0.190). ZBP1 association with mt-associated Z-DNA was more pronounced when directly compared with non-mt-associated Z-DNA. In PMA NETs, ZBP1 signal within mt-associated Z-DNA-positive regions was a median 2.68x higher than within non-mt-associated Z-DNA-positive regions (range 0.99–4.94x, n = 8; Wilcoxon p = 0.016), whereas the corresponding comparison following A23187 stimulation showed a median 3.17x enrichment (range 1.70–3.47x, n = 5, but p = 0.063). Importantly, ZBP1 signal within the mt-associated Z-DNA compartment was not elevated relative to mt-associated Z-DNA-negative regions (PMA median 0.90x, n = 8, p = 0.742; A23187 median 0.94x, n = 6, p = 1.00), indicating that the increased ZBP1 signal associated with mt-associated Z-DNA is specific relative to other Z-DNA-containing regions rather than reflecting a generalized increase in ZBP1 signal within mt-associated NET regions. Together, these findings support an association of ZBP1 with extracellular NET Z-DNA that is particularly pronounced within mt-associated Z-DNA.

**Figure 3:**
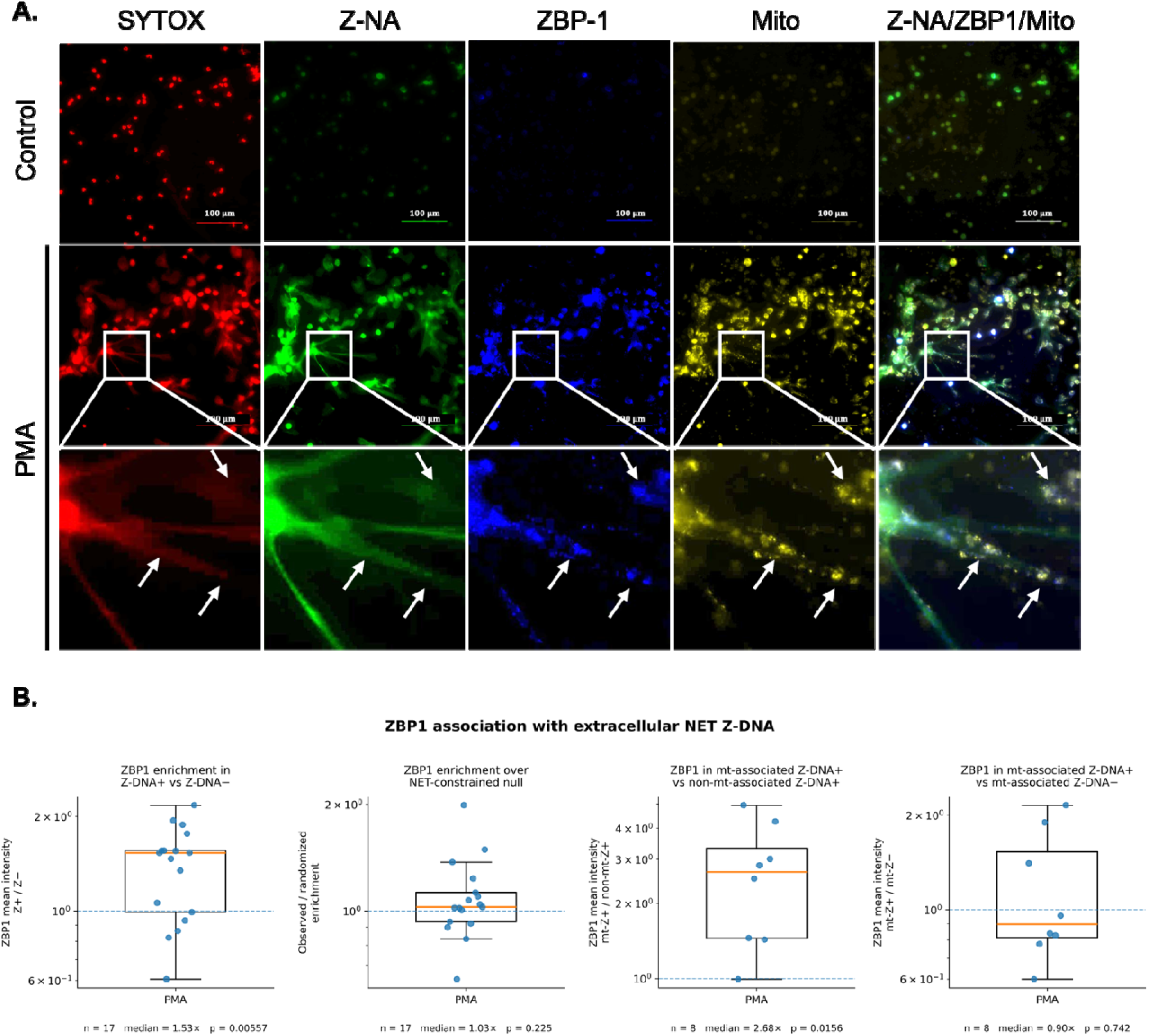
ZBP1 is associated with NETs Z-DNA with particular enrichment at mt-associated Z-DNA. **(A)** ZBP1 co-localization with mt-associated Z-DNA in NETs eDNA. Immunofluorescence images of neutrophils with and without PMA stimulation, stained for non-Z-DNA (SYTOX, Red), Z-DNA (Z-NA, green), ZBP1 (blue), and mitochondria (yellow), along with corresponding merged images. Insets (arrows) represent magnified views of the corresponding regions of interest (ROIs) from the panels above. **(B)** ZBP1 association with extracellular Z-DNA was assessed in PMA-induced NETs by comparing ZBP1 signal within Z-DNA-positive regions with corresponding non-Z-DNA regions and with NET-constrained randomized controls. Left, ZBP1 mean intensity was enriched in Z-DNA-positive (Z□) regions relative to Z-DNA-negative (Z □) regions (n = 17 images; median enrichment = 1.53-fold; p = 0.00557). Second, observed ZBP1 enrichment over a NET-constrained randomized null distribution was modest (n = 17; median enrichment = 1.03-fold; p = 0.225). Third, within Z-DNA-positive regions, ZBP1 signal was further enriched in mitochondria-associated Z-DNA-positive regions relative to non-mitochondria-associated Z-DNA-positive regions (n = 8; median enrichment = 2.68-fold; p = 0.0156). Right, comparison of ZBP1 signal within mitochondria-associated Z-DNA-positive and mitochondria-associated Z-DNA-negative regions showed no significant enrichment (n = 8; median enrichment = 0.90-fold; p = 0.742). Box plots show the median and interquartile range, with whiskers indicating the observed range; individual points represent analyzed images. Dashed horizontal lines indicate an enrichment value of 1.0 (no enrichment).

### Oxidative DNA damage is broadly associated with NET DNA and exhibits asymmetric localization within Z-DNA regions

Finally, given prior links between Z-DNA formation and oxidative DNA damage [15], we examined the relationship between Z-DNA and 8-oxoG. 8-oxoG was consistently detected within NETs DNA and exhibited non-random spatial association with both Z-DNA and non-Z-DNA (Supplementary Fig. S12). Colocalization analysis revealed pronounced asymmetry, with a substantial fraction of 8-oxoG localizing within Z-DNA (median Manders M2 = 0.37) while only a minority of Z-DNA overlapped with 8-oxoG (median M1 = 0.07; p = 0.027). To more directly quantify this relationship, 8-oxoG intensity within Z-DNA-positive DNA was compared to that in adjacent Z-DNA-negative NET DNA on a per-image basis: 8-oxoG was higher within Z-DNA-positive regions in 9 of 9 independently analyzed images, with a median enrichment of approximately 3.3-fold, robust across a range of Z-DNA positivity thresholds tested (median enrichment 2.4–3.6-fold across thresholds from 0.15–0.35). This association persisted after controlling for local DNA abundance, with Z-DNA and 8-oxoG signal remaining positively associated after partialling out local SYTOX intensity (median partial correlation ≈ 0.77). 8-oxoG signal was further enriched immediately proximal to Z-DNA, with intensity within 0–2 pixels of Z-DNA exceeding that at 5–10 pixels in 8 of 8 analyzable images (median near-to-far enrichment ≈ 3.6-fold), indicating that this spatial relationship extends beyond simple pixel-level overlap into the local environment surrounding Z-DNA suggesting preferential enrichment of 8-oxoG for Z-DNA over non-Z-DNA. Antioxidant treatment (vitamin C) suppressed NETosis globally, precluding direct assessment of its specific effect on Z-DNA formation within NETs.

### Z-DNA fraction of NETs persists nuclease-mediated degradation

*In vitro* and biofilm associated Z-DNA have been demonstrated to exhibit intrinsic resistance to nuclease-mediated degradation. Hence, we investigated whether this intrinsic property is preserved within NETs eDNA and can be implicated as a driver for overall NETs persistence to nucleases. To assess this, after stimulation, NETs were treated with DNase I and DNase1L3, two physiologically relevant nucleases involved in eDNA clearance, with DNase I targeting accessible eDNA and DNase1L3 being particularly effective against chromatin- or protein-associated DNA [49]. DNA integrity and Z-DNA localization were subsequently assessed by immunofluorescence analysis.

In PMA-induced NETs, extensive eDNA structures were readily detected, with Z-DNA signal distributed along filamentous DNA networks (Fig. 4). Following 15 minutes of nuclease treatment (DNase I or DNase1L3), the overall eDNA signal was reduced, indicating degradation of bulk DNA. However, Z-DNA–positive structures persisted under both DNase I and DNase1L3 conditions, remaining detectable within residual filamentous eDNA networks. Merged images revealed that regions retaining Z-DNA signal frequently corresponded to nuclease-resistant DNA fragments, whereas non-Z-DNA signal was substantially reduced, indicating that Z-DNA–containing regions exhibit reduced susceptibility to enzymatic degradation. We quantified these effects using our image analysis pipeline on a donor-sample matched dataset. We first generated baseline mean total Z-DNA skeleton length and total NET DNA intensity for each donor after PMA treatment and computed the fraction remaining after DNase1 and DNase1L3 treatments. This analysis revealed that both DNases treatments significantly altered NETs. There was a significant increase in overall DNA degradation (DNase1 treatment: median Δ = 0.572, p = 0.016), confirming enzymatic activity. However, the fraction of Z-DNA remaining was largely unchanged across DNase treatments (DNase1: median Δ = −0.642, p = 0.578; and DNase1L3: median Δ = 0.103, p = 0.844), indicating that Z-DNA is not proportionally lost during DNase-mediated degradation. These findings show that Z-DNA-containing regions are preferentially retained within NET structures following physiologically relevant nuclease challenges, consistent with a potential contribution to NET persistence.

**Figure 4:**
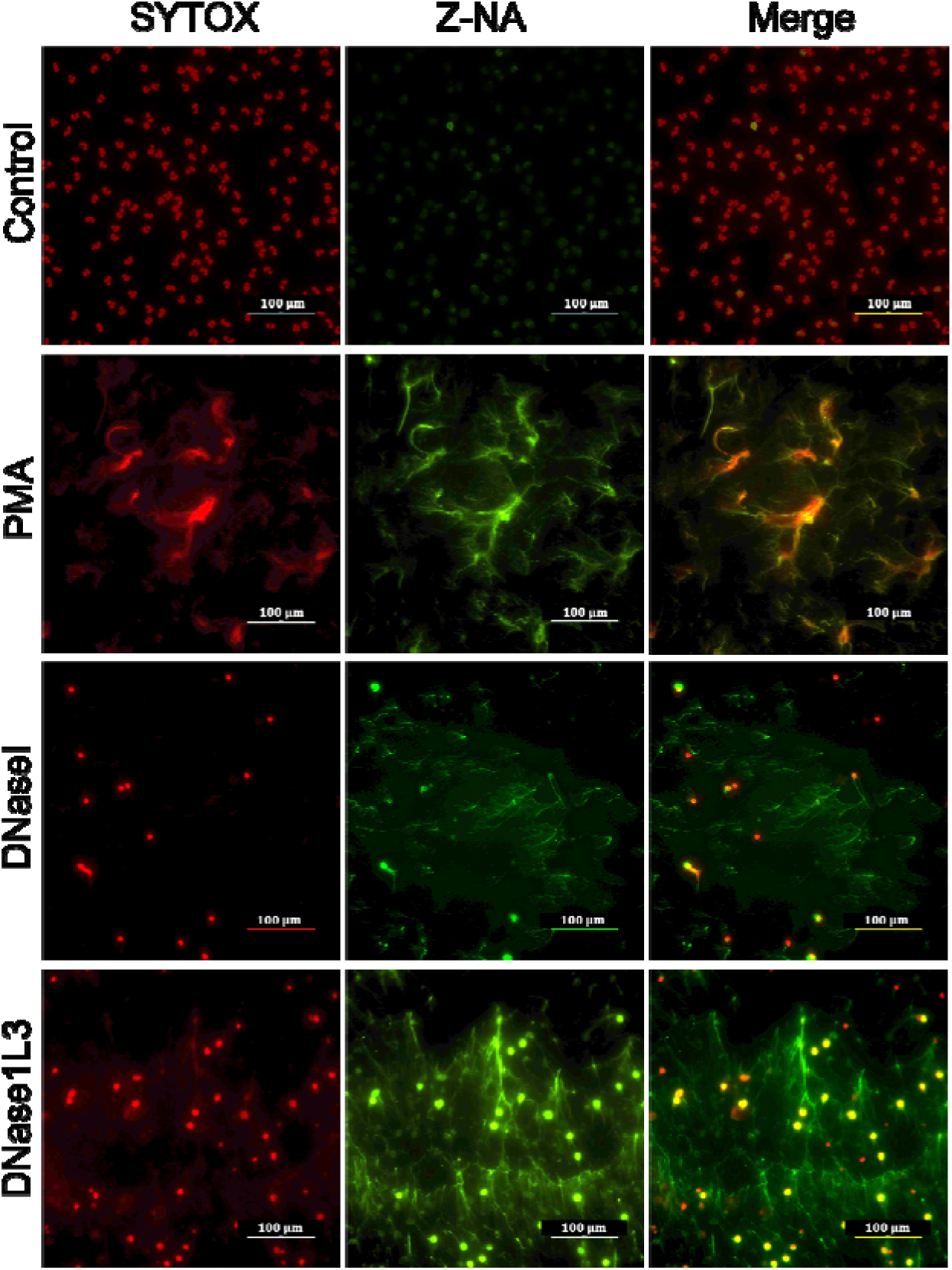
NETs Z-DNA is preferentially retained after DNase digestion. Representative immunofluorescence images of neutrophils stimulated with PMA (1 µM, 4 h) to induce NETosi and subsequently treated with DNase I or DNase1L3 under the indicated conditions. Non-Z-DNA was stained with SYTOX (red), and Z-DNA was detected using the Z22 anti–Z-DNA antibody (green). Merged images show the localization of Z-DNA within total DNA structures. Scale bars, 100 μm.

### Modulation of DNA conformation restores nuclease susceptibility of NET-derived DNA

We next investigated whether altering DNA conformation could restore susceptibility to enzymatic degradation. For this, neutrophils were treated with chloroquine (CQ), a small molecule known to stabilize B-DNA and disfavor Z-DNA formation and previously shown to convert Z to B-DNA in biofilms, during the final hour of NETosis. After that, we performed immunofluorescence analysis of NET structures alone or in combination with DNase.

PMA stimulation induced robust NET formation characterized by extensive eDNA structures with prominent Z-DNA signal. Upon CQ treatment, a marked reduction in Z-DNA signal was observed, accompanied by a relative increase in non-Z-DNA signal across all 3 donor-matched experiments corresponding to a 75% reduction in the geometric mean Z/non-Z ratio (CQ/PMA = 0.255; 95% CI, 0.040–1.65), indicating a shift in DNA conformation in Z-DNA to non-Z-DNA. Combined CQ and DNase treatment led to near-complete loss of eDNA structures, consistent with increased nuclease susceptibility accompanying the CQ-associated shift in DNA conformational state (Fig 5A).

**Figure 5:**
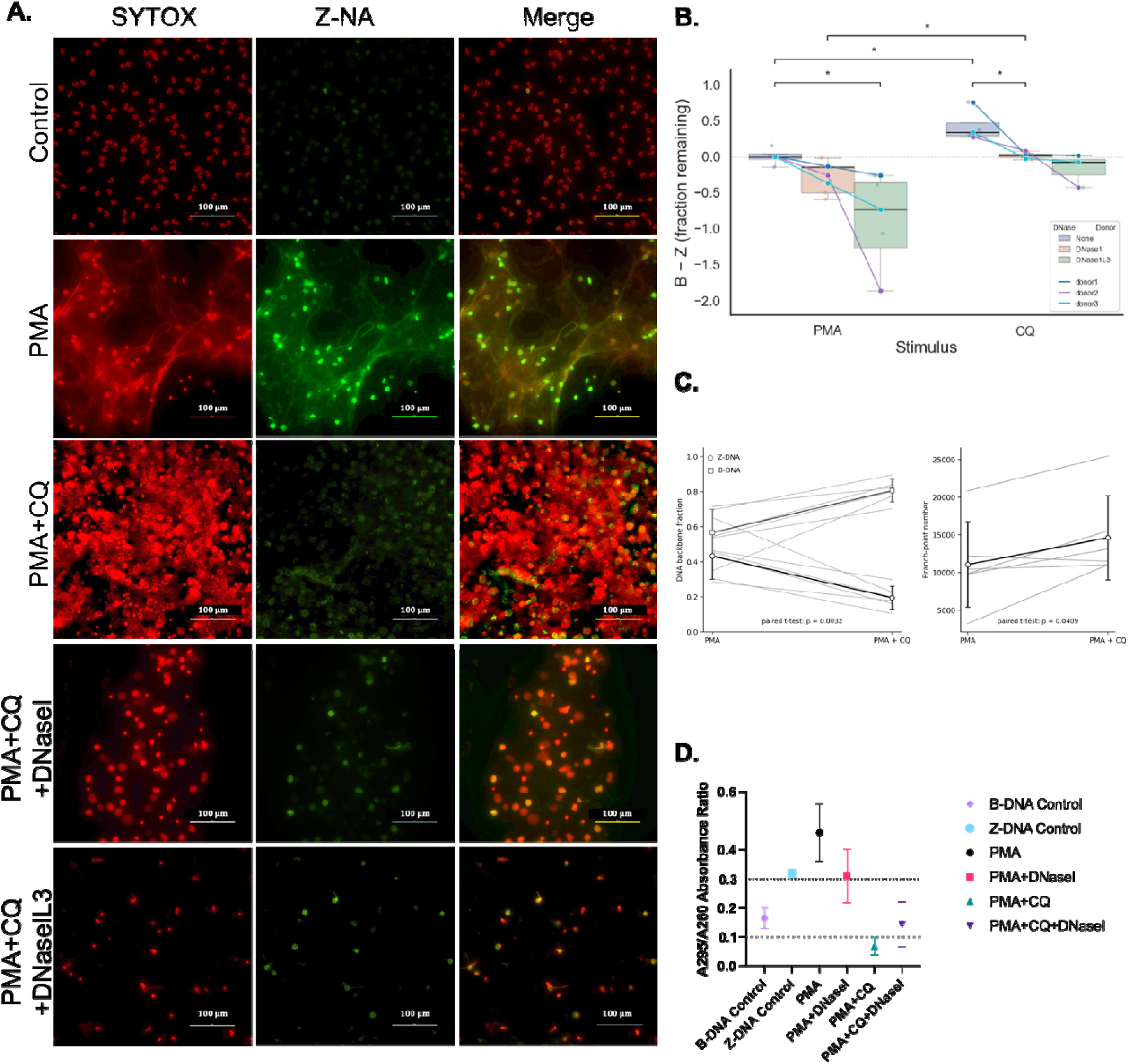
CQ reduces Z-DNA in NETs and increases susceptibility to nucleases. **(A)** Representative immunofluorescence images of neutrophils following PMA-induced NETosis with or without CQ and nuclease treatment. Non-Z-DNA was visualized using SYTOX Red, and Z-DNA was detected using Z22 antibody. Conditions shown include PMA, PMA + CQ, and combined CQ and nuclease treatments. Scale bars, 100 µm. **(B)** Sample-matched analysis of Z- and non-Z-DNA total skeleton length retained after CQ or nuclease treatments compared to baseline PMA or PMA+CQ levels on 24 images collected from 3 donors. The fractions of both DNA conformations retained after treatments were compared across conditions as the difference between non-Z- and Z-DNA (non-Z − Z) of the remaining fraction. Statistical analysis was performed using the Wilcoxon signed-rank test. Significant reversal in Z-DNA to non-Z-DNA in NETs upon CQ treatment and subsequent reduction in the converted non-Z-DNA with nuclease treatment. Same trend observed with both nucleases, but the assay did not reach significance with DNase1L3 after CQ treatment. (**C)** Donor-matched NETs were analyzed before and after CQ treatment. Left, Z-DNA (circles) and non-Z-DNA (squares) backbone fractions in PMA- and CQ-treated NETs. Individual gray lines connect donor-matched measurements, with black symbols and lines showing the mean ± SD. CQ treatment significantly decreased the Z-DNA backbone fraction and increased the non-Z-DNA backbone fraction (paired t-test, p = 0.0032). Right, branch-point number in donor-matched PMA and PMA + CQ NETs. Individual gray lines represent paired donors and black symbols and lines show the mean ± SD. CQ treatment was associated with an increase in branch-point number (paired t-test, p = 0.0409) **(D)** Absorbance-based analysis of NET-derived DNA and control oligonucleotides. Z-DNA and B-DNA oligonucleotides were included as references. Absorbance values were measured for DNA isolated from the indicated treatment conditions at 295nm and 260nm and the ratios were computed.

We quantified these observations the same way as above, by performing a donor-sample matched analysis. However, since this time we were estimating the potential shift of Z-DNA to B-DNA by CQ in NETs, we also computed fraction of non-Z-DNA total skeleton length after DNase CQ and DNase treatment per donor relative to the PMA and CQ baselines respectively. We then estimated the difference between the non-Z-DNA fraction remaining and Z fraction remaining (non-Z_minus_Z) to assess the relative composition of non-Z and Z-DNA in the NETs after treatment relative to the baseline. CQ stimulation shifted NET DNA toward a more non-Z-DNA–enriched state relative to Z-DNA in donor-matched samples, as reflected by an increase in non-Z_minus_Z (Δ = 0.33, p = 0.031) compared to PMA (Fig. 5B). Notably, this non-Z-DNA enriched state was attenuated following DNase treatments under CQ conditions (median Δ = −0.361, p = 0.031 for DNase1, n = 6; median Δ = −0.574, p = 0.125, n = 4, for DNase1L3), indicating preferential loss of non-Z-DNA relative to Z-DNA. The DNase1L3 comparison did not reach significance; given the observed effect size and donor-to-donor variability, our data cannot distinguish between a true absence of effect and insufficient sample size to detect an effect of this magnitude. Together, these findings support a model in which CQ promotes a non-Z-DNA–favored structural state within NETs that remains dynamically susceptible to nuclease-mediated remodeling.

To determine whether the Z-to-non-Z shift was accompanied by changes in the spatial organization of the NET scaffold, we first assessed global NET architecture. CQ treatment produced a pronounced conformational shift in the NET-associated DNA scaffold, with the Z-DNA backbone fraction decreasing and the non-Z-DNA backbone fraction increasing in donor-matched samples (both p = 0.0032; Fig. 5C). Z-DNA mean intensity and total Z-DNA intensity were also reduced following CQ treatment (p = 0.0043 and p = 0.0080, respectively), providing independent evidence for the conformational shift. CQ treatment did not result in collapse of the NET scaffold; instead, several architectural measures showed directional increases, including NET area (∼40%), total network length, mesh spacing (∼122%), and SYTOX skeleton content (∼49%). Among the topology measures examined, branch-point number increased in donor-matched samples (p = 0.0409; Fig. 5C), consistent with remodeling of network connectivity accompanying the conformational shift. Because multiple architectural metrics were evaluated and the branch-point association did not remain significant after correction for multiple comparisons, we interpret this finding as evidence of an associated topological change rather than broad, statistically significant remodeling across the NET architecture. Together, these data indicate that CQ-mediated Z-to-B conversion is accompanied by modest remodeling of NET topology without the collapse of the existing extracellular scaffold.

To independently validate this structural shift, we analyzed NET-derived DNA using absorbance-ratio-based measurements. Based on established spectral characteristics, higher A295/A260 ratios (≥0.3) are consistent with Z-DNA–like conformations, whereas lower values are associated with B-DNA–like structures [24]. Compatible with this, Z-DNA oligonucleotide controls, as well as DNA isolated from PMA- and PMA + DNase I–treated samples, exhibited absorbance values at or above this threshold, consistent with Z-DNA–enriched conformations. In contrast, B-DNA oligonucleotide controls and CQ-treated samples, including PMA + CQ and PMA + CQ + DNase I conditions, showed reduced absorbance values, consistent with a shift toward non-Z-DNA–like conformations (Fig. 5D). It is important to note that CQ is a pleiotropic agent with additional activities beyond DNA conformational modulation, including TLR9 antagonism, inhibition of lysosomal/autophagic processes, and direct DNA intercalation. As CQ was present during the final hour of active NETosis, we cannot exclude the possibility that these alternative mechanisms contribute to the observed restoration of nuclease susceptibility. However, the concordant shift in absorbance-ratio conformational signatures (Fig. 5D) supports a genuine contribution of Z-to-non-Z DNA structural change to this phenotype.

Our orthogonal lines of evidence support the presence of Z-DNA in NETs and its preferential retention after nuclease challenge. CQ treatment shifted NET DNA toward a non-Z state and was accompanied by increased nuclease susceptibility; because CQ is pleiotropic, these experiments support an association between conformational state and susceptibility rather than attributing the phenotype exclusively to Z-DNA.

### Z:non-Z-DNA conformation is associated with coordinated NET architecture

The temporal analysis earlier established how Z-DNA accumulates within a developing NET scaffold, but it does not determine whether variation in the relative Z:non-Z-DNA composition itself is associated with a particular architectural state. We therefore asked whether NET architecture varied continuously with the measured Z:non-Z-DNA conformational state across independent perturbations, rather than along a single temporal trajectory. We therefore combined images from the PMA/CQ, CaCl□, and spermidine perturbation series and examined the relationship between the continuous log10-transformed Z:non-Z-DNA skeleton ratio and NET architecture. The analysis included 53 images with complete measurements: 19 from the PMA/CQ series, 15 from the CaCl series, and 19 from the spermidine series. Across the combined dataset, increasing Z:non-Z-DNA skeleton ratio was associated with a coordinated shift in overall NET architecture. Principal component analysis of 26 independent architecture metrics showed that PC1 explained 46.95% of the total architectural variance and was negatively associated with Z:non-Z-DNA ratio (Spearman ρ = −0.610, p = 1.27 × 10□□; Fig. 6A). At the individual-metric level, increasing Z:non-Z-DNA ratio was associated with greater filament thickness (ρ = 0.719, p = 1.35 × 10□□), mesh spacing (ρ = 0.674, p = 3.27 × 10□□), and average branch length (ρ = 0.489, p = 2.04 × 10□□), and with lower filament density (ρ = −0.711, p = 2.51 × 10□□), junction density (ρ = −0.770, p = 1.52 × 10□¹¹), and triple-point frequency (ρ = −0.779, p = 6.48 × 10□¹²) (Fig. 6B). NET area (ρ = −0.325, p = 0.0177), pore count (ρ = −0.293, p = 0.033), and mean pore area (ρ = −0.344, p = 0.0117) also showed nominal pooled associations, but these did not remain significant after adjustment for perturbation arm and multiple testing. To determine whether the observed relationships were independent of the experimental perturbation series, we performed arm-adjusted linear models across all 26-architecture metrics, with log10□(Z:non-Z-DNA skeleton ratio) as the continuous predictor and perturbation arm as a covariate. After Benjamini–Hochberg correction, Z:non-Z-DNA ratio remained significantly associated with filament density (β = −20.5, q = 8.49 × 10□□), thickness (β = 109.6, q = 2.65 × 10□□), SYTOX skeleton content (β = −1.23 × 10□□, q = 0.00107), branch-point number (β = −8.23 × 10³, q = 0.00107), total network length (β = −1.23 × 10□□, q = 0.00107), triple-point frequency (β = −0.00396, q = 0.00174), junction density (β = −0.00577, q = 0.00445), average branch length (β = 11.9, q = 0.00469), and mesh spacing (β = 100.5, q = 0.00903), with additional FDR-significant associations for orientation index (β = 0.1404, q = 0.0209), quadruple-point frequency (β = −0.001803, q = 0.0213), dispersion index (β = −20.66, q = 0.0213), maximum filament length (β = −374.3, q = 0.0213), and non-Z-DNA fragmentation index (β = 0.002634, q = 0.0231). Therefore, these data demonstrate that the Z:non-Z-DNA conformational state is continuously associated with coordinated changes in higher-order NET architecture across independent perturbation series, with increasing Z:non-Z-DNA ratio associated most consistently with thicker, more widely spaced filaments and longer branches, together with reduced filament density and junctional subdivision. Because this analysis examines covariance across perturbation-induced changes in the Z:non-Z-DNA axis, it establishes a reproducible association between DNA conformation and NET architecture where a higher Z:non-Z-DNA ratios corresponded to a coarser mesh composed of thicker, more widely spaced DNA elements with longer structural segments and fewer points of junctional subdivision, rather than simply a greater amount of eDNA.

**Figure 6:**
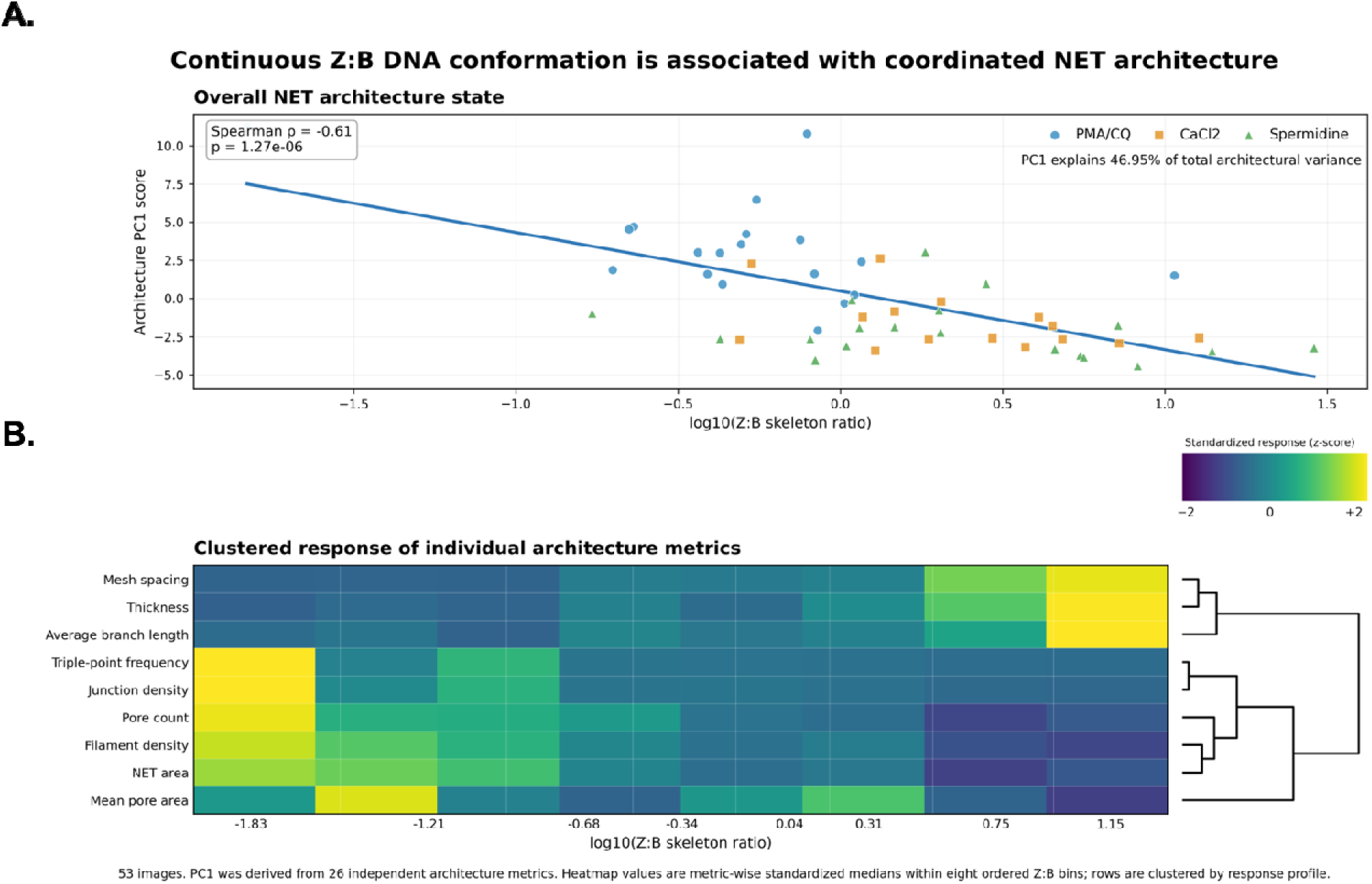
Z:non-Z-DNA conformation is associated with coordinated NET architecture. **(A)** Relationship between continuous Z:non-Z-DNA conformation and overall NET architectural state across 53 images from the PMA/CQ, CaCl□, and spermidine perturbation series. The x-axis shows the log10-transformed Z:non-Z skeleton ratio, and the y-axis shows the first principal component (PC1) derived from 26 independent NET architecture metrics. PC1 accounted for 46.95% of total architectural variance. Each point represents an individual image, with colors/symbols indicating perturbation series. The pooled association between Z:non-Z ratio and PC1 was assessed by two-sided Spearman rank correlation (ρ = −0.610, p = 1.27 × 10□□); the line represents an ordinary least-squares fit for visualization. **(B)** Heatmap showing the respons of representative NET architectural metrics across eight ordered bins of the Z:non-Z skeleton ratio. Values represent metric-wise standardized median responses within each Z:non-Z bin; rows were hierarchically clustered according to their response profiles. Increasing Z:non-Z ratio wa associated with increased filament thickness, mesh spacing, and average branch length and with decreased filament density, junction density, and triple-point frequency. The continuous associations were evaluated across all 53 images by Spearman rank correlation and independently tested using linear models adjusted for perturbation arm, with Benjamini–Hochberg correction across the 26-metric screen. Metrics including filament density, thickness, SYTOX skeleton content, branch-point number, total network length, triple-point frequency, junction density, average branch length, and mesh spacing remained significantly associated with Z:non-Z ratio after adjustment and FDR correction.

### Z-DNA fraction of NETs is immunostimulatory and induces type I interferon responses

Given the established role of NETs in activating innate immune pathways and promoting type I interferon responses, we next asked whether Z-DNA-enriched NET fractions are associated with enhanced immunostimulatory activity. To isolate conformationally distinct NET-derived DNA fractions, NETs generated from PMA-stimulated healthy donor neutrophils were immunoprecipitated using the Z-DNA–specific Z22 antibody, yielding Z-DNA–enriched and corresponding non-Z-DNA fractions. Healthy donor PBMCs were then stimulated in a dose-dependent manner with either 2 µg or 5 µg of each fraction, and IFN-α production was quantified by ELISA. LPS stimulation was included as an inflammatory positive control, and cells-only wells were used for background normalization.

Z-DNA–enriched fractions induced greater IFN-α production from PBMCs compared to non-Z-DNA fractions across both doses (Fig. 7). The data suggest a dose-responsive pattern, with the 5 µg Z-DNA condition yielding a notably higher median fold-change (2.3×) relative to non-Z-DNA compared to the 2 µg condition (1.6×). To analyze immunostimulatory potential in an input-concentration-agnostic manner, data were pooled across both doses, revealing that Z-DNA–enriched fractions consistently induced higher IFN-α than non-Z-DNA fractions (Wilcoxon p = 0.001). Notably, this effect was donor-independent: every individual donor across both doses exhibited a fold-change greater than 1 for Z-DNA enriched versus non-Z-DNA fractions stimulations (median fold-change 1.80, IQR 1.41–2.59; Wilcoxon W = 0, p = 0.001), indicating a robust and reproducible immunostimulatory signal associated with the Z-DNA–enriched fraction. While these data show that Z-DNA-enriched NET fractions are associated with enhanced type I interferon induction in human PBMCs at matched eDNA concentrations, the effects cannot be attributed solely to Z-DNA because the immunoprecipitated fractions can differ in associated proteins and other molecular components.

**Figure 7:**
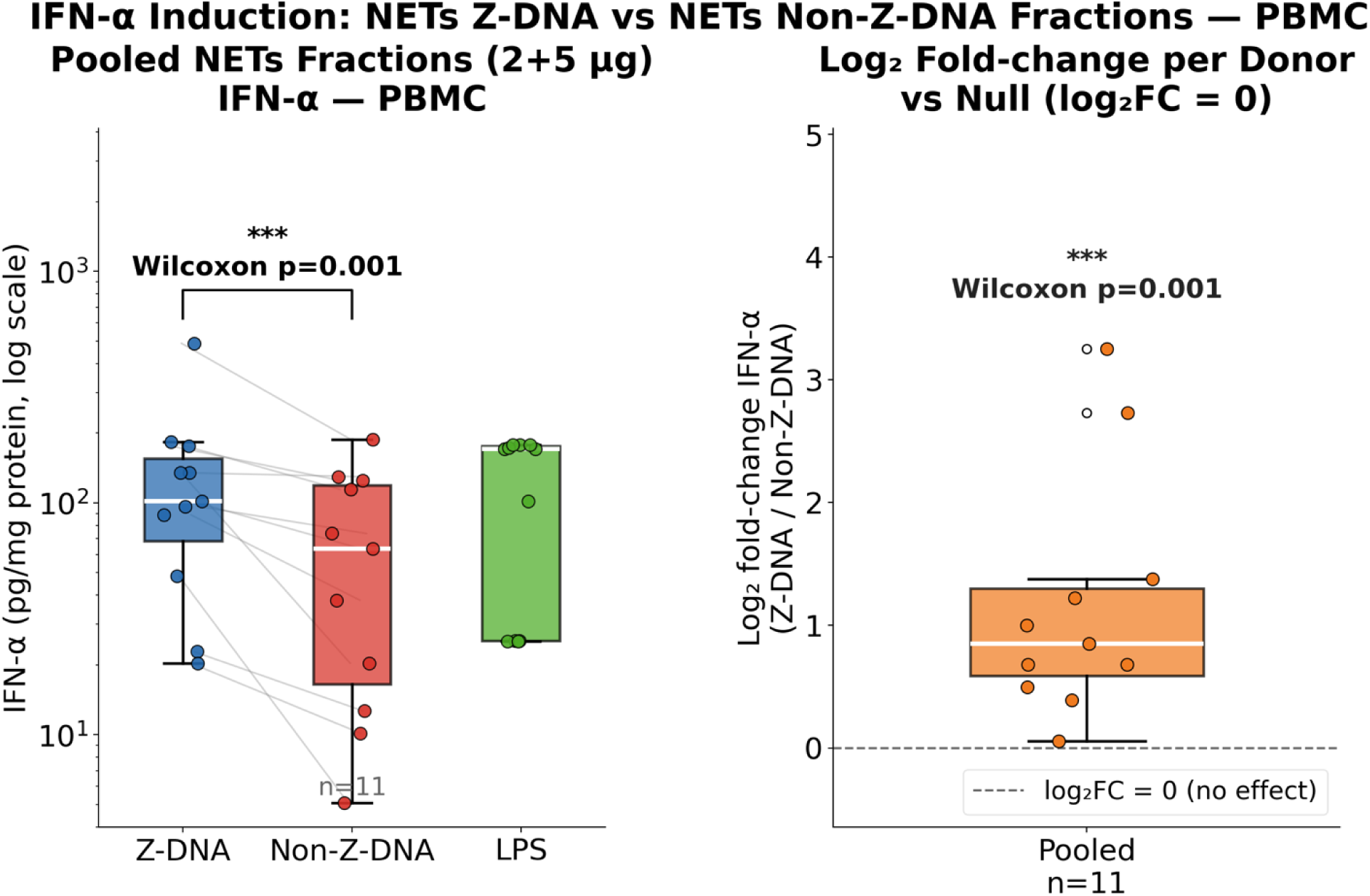
NETs Z-DNA fraction induces greater IFN-α in human PBMCs. PBMCs stimulated with NETs Z-DNA or Non-Z-DNA fractions (n=11 paired observations, 3 donors). IFN-α (pg/ml protein) shown with LPS as positive control (left); fold change/donor relative to FC=1 null (right). Z-DNA consistently induced greater IFN-α across all donors (median FC=1.80, IQR 1.41–2.59). Wilcoxon signed-rank test; p=0.001.

**Figure 8:**
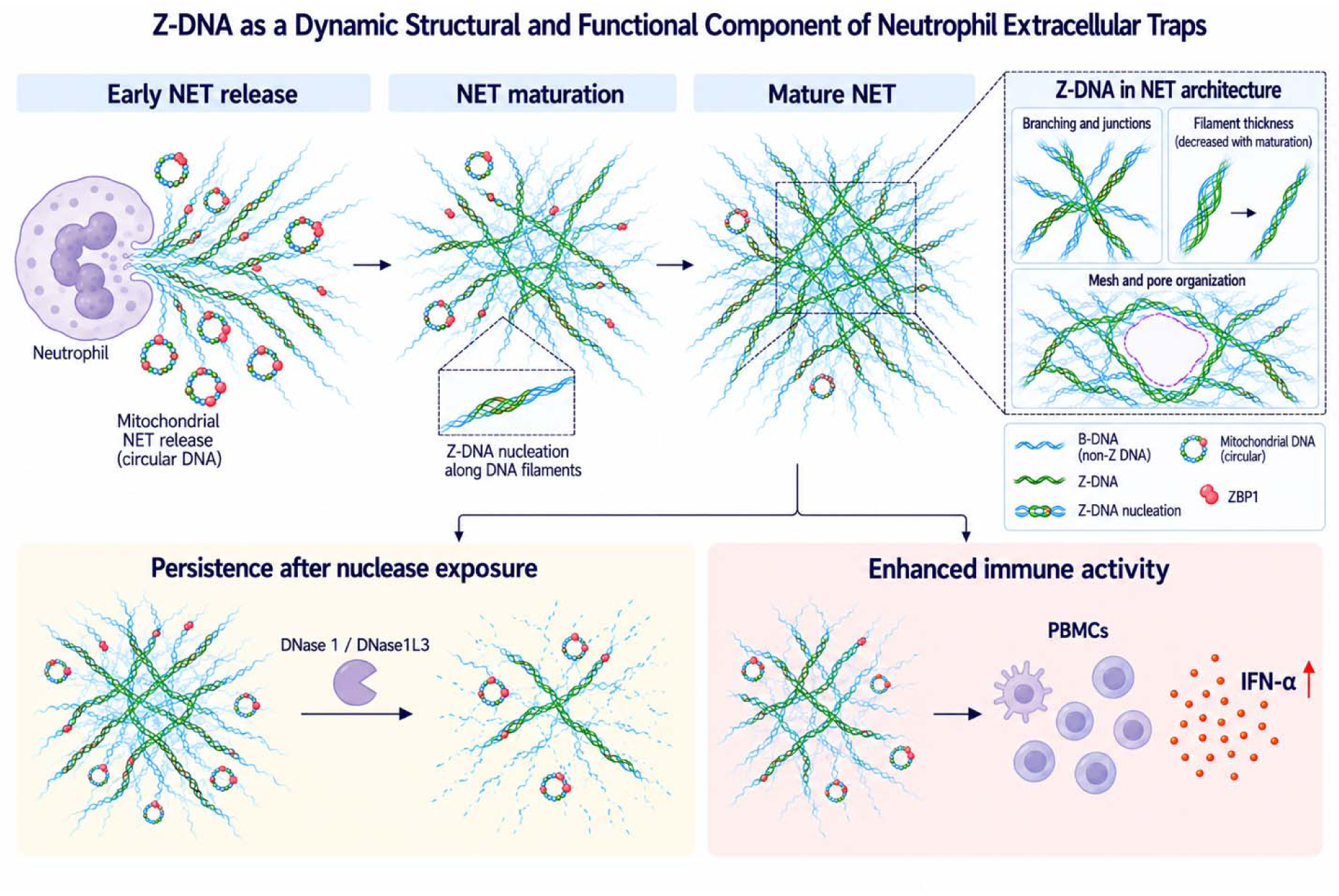
Z-DNA as a dynamic structural and functional component of NETs. Schematic depicting the proposed integration and functional consequences of Z-DNA within the NET scaffold. During early NET release, mitochondrial and oxidative cues may contribute to the appearance of spatially discrete Z-DNA domains, which progressively accumulate within the expanding extracellular network during NET maturation. Z-DNA remains associated with features of NET architecture, including branching, junctional complexity, filament geometry, and pore organization. ZBP1 shows preferential association with mt-associated Z-DNA, whereas global ZBP1/Z-DNA association elsewhere in the NET scaffold is not greater than expected from NET geometry alone. Following nuclease challenge, Z-DNA is preferentially retained within the residual NET structure. Z-DNA–enriched NET fractions also exhibit enhanced immunostimulatory activity in PBMCs, reflected by increased IFN-α production. Together, these observations support a model in which Z-DNA represents a dynamic component of the NET scaffold whose abundance and spatial distribution are associated with its organization, preferential persistence, and biological activity.

## Discussion

Our findings identify Z-DNA as an intrinsic component of eDNA within NETs, extending our previous demonstration that NET DNA can adopt noncanonical conformations such as G4s with catalytic DNAzyme activity [5]. Z-DNA was detected across diverse NETosis inducing stimuli and pathways, including *in vitro* suicidal and vital NETosis and *ex vivo* models. The consistency in Z-DNA formation across diverse inflammatory triggers (Fig. 1B and Supplementary Fig. S5) indicates that Z-DNA is not stimulus-specific but rather represents a convergent structural outcome of eDNA exposed to the physicochemical conditions of the NET milieu. Z-form DNA has recently been reported in granulomatous inflammatory lesions in tuberculosis and sarcoidosis patient tissue [50] and has been linked to exogenous DNA-remodeling, structural integrity, and persistence in microbial biofilms [7, 8, 16]. In fact, bacterial DNABII proteins can remodel eDNA and promote Z-DNA formation in NETs. Our findings extend these observations by demonstrating that host-derived NET DNA can adopt the Z conformation intrinsically, without requiring bacterial or other exogenous remodeling, establishing Z-DNA as a bona fide component of the NET extracellular matrix.

A fundamental aspect of this study is the delineation of a temporal framework for Z-DNA accumulation during the progression of NETosis. Z-DNA continues to accumulate throughout NET maturation primarily through increasing numbers of spatially discrete domains rather than simple elongation of pre-existing domains. Of note, because these measurements derive from fixed, diffraction-limited images, they do not directly resolve molecular nucleation events. However, the progressive increase in discrete Z-DNA domain number, together with the relative plateau in mean domain length, is suggestive of progressive nucleation of additional Z-DNA domains during NET maturation (Fig. 2C). This suggests that DNA extrusion does not represent the endpoint of NET DNA organization; instead, the extracellular scaffold remains structurally dynamic after release. Consistent with this, progressive nucleosome stripping during NET maturation exposes naked DNA at which the energetic barrier to Z-DNA formation is lowest, providing a structural basis for the domain-number-driven expansion we observe [4]. The enrichment of Z-DNA with spermidine and CaCl further supports sensitivity of this conformational state to the extracellular physicochemical environment. These findings suggest that the changing ionic, oxidative, and mechanical environment of the maturing NET can continually reshape the conformational state of eDNA.

The preferential association of Z-DNA with branching, junctional complexity, and connectivity indicates that Z-DNA accumulation accompanies remodeling of the higher-order NET scaffold. Importantly, absolute Z-DNA abundance and relative Z:non-Z-DNA composition represent distinct dimensions of NET organization: during PMA-induced maturation, Z-DNA increased substantially without a monotonic increase in the overall Z:non-Z-DNA ratio, becoming increasingly incorporated into a network that expanded, thinned, branched, and developed greater junctional connectivity. The concurrent increase in pore number without a change in mean pore area further indicates progressive network elaboration and subdivision. Such remodeling could increase the number of discrete sites available for physical retention of extracellular material without substantially altering the dimensions of individual pores [4]. In contrast, across independent PMA/CQ, CaCl□, and spermidine perturbations, increasing Z:non-Z-DNA ratio was associated with a distinct architectural state characterized by thicker, more widely spaced filaments, longer branches, and reduced filament and junctional density, with the global architectural axis explaining 46.95% of variance (Spearman ρ = −0.610, p = 1.27 × 10□□; Fig. 6). Thus, Z-DNA accumulation during maturation and variation in relative Z:non-Z-DNA composition describe complementary, non-equivalent aspects of NET architecture rather than a single linear trajectory.

The spatial and temporal distribution of Z-DNA further suggests that its formation is influenced by the molecular origin and local environment of NET DNA. Mt-associated Z-DNA was enriched early during NETosis before the progressive accumulation of nuclear-associated Z-DNA, while ZBP1 showed preferential association with the mt-associated Z-DNA compartment despite the absence of significant global ZBP1 enrichment across the NET Z-DNA network. This pattern argues against uniform recognition of extracellular Z-DNA and instead suggests that DNA origin, accessibility, or local molecular context may influence ZBP1 engagement. The broader literature establishes ZBP1 as a sensor of Z-conformation nucleic acids, supporting the biological plausibility of this interpretation [20, 51–56]. Additionally, the lack of reciprocity in global Z-DNA to ZBP1 engagement raises the possibility of the existence of other stabilizing proteins in the NETs milieu – like the Z domain containing ADAR1 or stabilization by the DNABII functional homolog HMGB1, consistent with the Human Protein Atlas reporting ADAR expression in neutrophils. The pronounced spatial association between 8-oxoG and Z-DNA provides an additional, albeit limited, link between the oxidative environment of NET formation and local Z-DNA organization. Our data, therefore, suggest that Z-DNA formation is shaped not only by the global extracellular environment but also by the biochemical history and local context of individual DNA domains.

The preferential retention of Z-DNA-containing regions following DNase I and DNase1L3 treatment indicates that DNA conformational state may be associated with persistence of NET eDNA, complementing established mechanisms involving DNA accessibility and associated proteins [2, 57]. Consistent with this possibility, shifting the Z:B equilibrium toward a non-Z-DNA state with CQ reduced preferential Z-DNA retention and increased susceptibility to nuclease degradation (Fig. 5), although the pleiotropic effects of CQ preclude attributing this effect exclusively to DNA conformation. The potential functional consequences of this persistence extend beyond structural integrity. Z-DNA-enriched NET fractions induced greater IFN-α production than non-Z-DNA fractions at matched eDNA concentrations, associating Z-DNA enrichment with increased immunostimulatory activity. This finding is consistent with previous work showing that NET-associated oxidized mitochondrial DNA can be strongly interferogenic and raises the possibility that the conformational state of persistent NET DNA contributes to how eDNA is sensed by immune cells. These findings raise the possibility that DNA conformational state contributes to the immunogenic properties of extracellular self-DNA, beyond DNA abundance, sequence, oxidation state, or subcellular origin; however, this interpretation remains associative in the present fractionation experiment. However, because the Z-DNA-enriched fractions also contain associated proteins and other molecular components, the fractionation experiment does not establish Z-DNA as the sole mediator of the IFN-α response, nor does it establish ZBP1 as the responsible sensor. Taken together, these findings link Z-DNA enrichment with two functionally relevant properties of NET eDNA—persistence following nuclease exposure and differential immunostimulatory activity—supporting a model in which DNA conformation may influence not only the physical longevity of the NET scaffold but also the biological consequences of the DNA that remains.

While our findings identify Z-DNA as an intrinsic component of NETs, certain aspects of the study should be interpreted within the context of current technical and experimental limitations. Quantitative imaging analyses incorporated multiple fields per donor and experimental condition, enabling high-resolution sampling of NET architecture across the observed systems. Because these measurements are hierarchically structured within donors and experiments, image-level statistical significance is interpreted as evidence for reproducible associations within the sampled datasets rather than as independent biological replication. Where relevant, donor-level analyses are used to assess the consistency and generalizability of these relationships. Although our *ex vivo* assays confirmed the presence of Z-DNA in NET-derived eDNA under more physiologically relevant extracellular conditions, reduced NET yield and increased background signal limited direct quantitative comparison with the *in vitro* datasets. Validation in patient-derived inflammatory specimens will therefore be an important next step; however, such analyses are complicated by high extracellular background signals, serum-associated nucleases and carrier proteins, and the fact that extracellular trap formation in whole blood or inflamed tissues is not exclusively neutrophil-derived, making cell-type-specific attribution of Z-DNA–positive structures challenging. Detection of DNA conformation in complex extracellular matrices also remains technically challenging. Commercially available reagents capable of selectively labeling canonical B-DNA without cross-recognizing other dsDNA conformations remain limited. We therefore used SYTOX Red operationally as a non-Z-DNA–enriched signal rather than as a definitive B-DNA marker, supported by evidence that this dye exhibits markedly reduced fluorescence upon interaction with Z-DNA. However, partial dye retention at mixed B/Z-DNA regions or structural junctions may lead to underestimation of the total Z-DNA burden. All imaging-based analyses were performed on two-dimensional widefield fluorescence micrographs, and the topology, connectivity, and spatial enrichment metrics reported here therefore reflect projected network architecture rather than the full three-dimensional organization of NETs in situ; future studies using confocal z-stack reconstruction or super-resolution approaches will be needed to resolve the three-dimensional distribution of Z-DNA within the NET scaffold and confirm whether the enrichment observed here persists across the depth of the structure. In addition, all experiments were performed using neutrophils from healthy donors, and whether the Z-DNA dynamics described here are altered in disease contexts such as SLE, where NET composition, immune complex formation, and eDNA clearance are fundamentally perturbed, remains to be determined. Finally, the exact contribution of Z-DNA itself, independent of local DNA packing, associated proteins, and the broader physicochemical effects of the perturbations used here, could not be directly assessed in the current study. Likewise, MitoTracker proximity identifies mt-associated Z-DNA but does not by itself establish mitochondrial sequence origin, and the Z-DNA-enriched PBMC-stimulation fractions may differ in associated molecular components. Broader functional consequences, including potential contributions to antimicrobial activity, pathogen trapping, and other effector functions, therefore remain to be established. This limitation reflects the absence of selective Z-DNA–specific inhibitors or structural modulators that do not concomitantly perturb intersecting inflammatory pathways and represents a key priority for future tool development.

We propose that NET DNA exists within a dynamic conformational continuum in which early mitochondrial-associated and oxidative cues may accompany the appearance of spatially discrete Z-DNA domains, followed by their continued accumulation within the extracellular scaffold. Variation in the Z:non-Z conformational state is associated with systematic changes in filament geometry and network connectivity, while Z-DNA-containing regions are preferentially retained following nuclease challenge and Z-DNA-enriched NET fractions show enhanced immunostimulatory activity. Together, these findings support a model in which Z-DNA is not an incidental conformation but a dynamic component of the NET scaffold associated with its organization, persistence, and biological activity; direct causal contributions of Z-DNA itself remain to be established.

## Author Contributions

Conceptualization: T.D.D, N.R.M. and S.Y.; Methodology: T.D.D., N.R.M. and S.Y.; Investigation: T.D.D. and N.R.M.; Visualization: T.D.D; Funding acquisition: S.Y.; Project administration: S.Y.; Supervision: S.Y.; Writing—original draft: T.D.D, N.R.M. and S.Y.; Writing—review & editing: T.D.D, N.R.M. and S.Y.

## Supplementary Data

Supplementary Data are available online.

## Data Availability

All data supporting the findings of this study are provided within the article and its Supplementary Data. Code used to analyze images and extract the metrics reported in this study are available at (https://github.com/nikhilram/net-dna-structure-analysis)

