## Supplementary Figures for "Z-DNA Reveals a Conformational Dimension of Neutrophil Extracellular Trap Organization"

**Supplementary Figure 1**

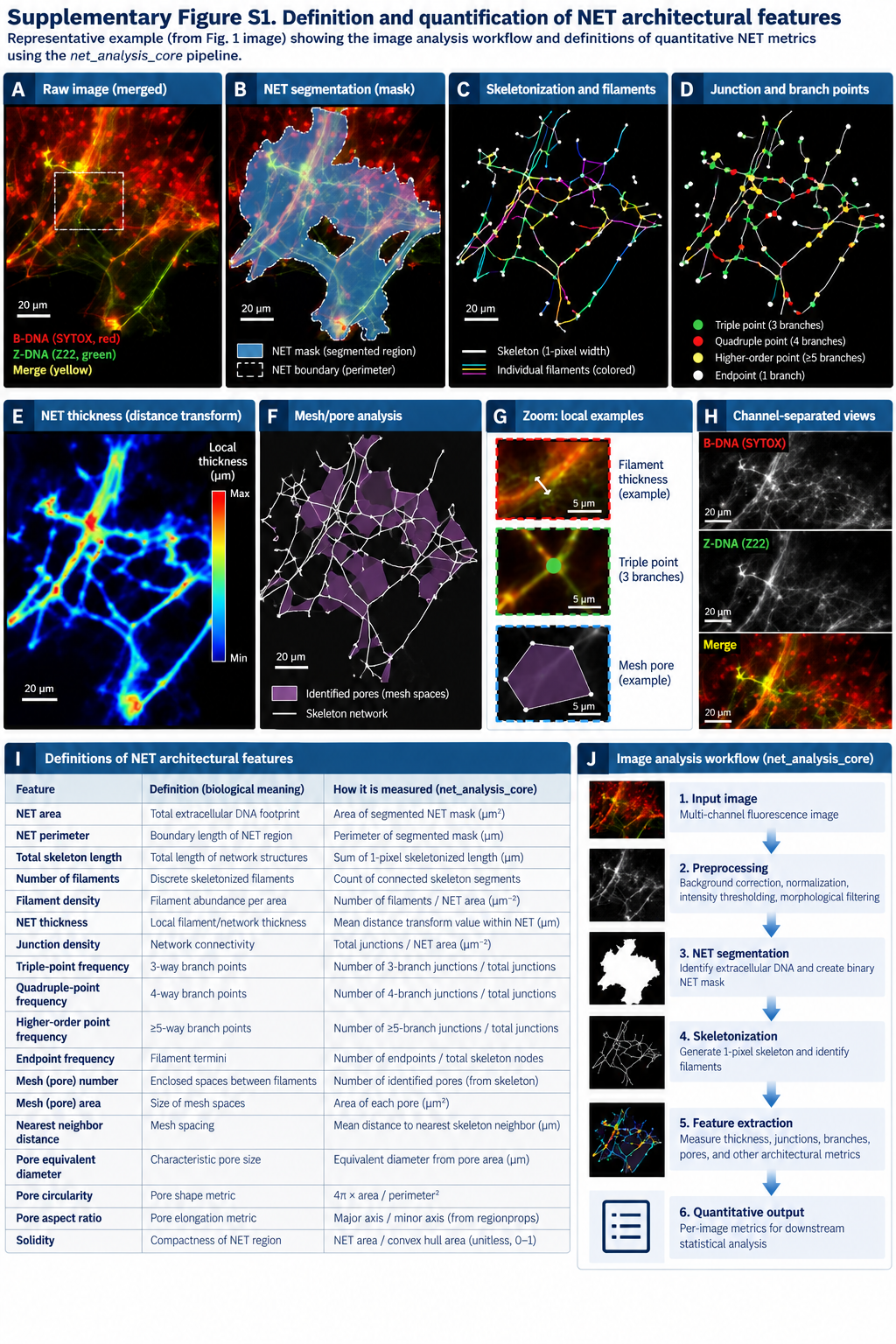

**Supplementary Figure S1: Image-analysis workflow for quantitative characterization of NET architecture and DNA composition. (A)** Representative multichannel NET image showing B-DNA (SYTOX, red), Z-DNA (Z22, green), and merged signal. **(B)** Segmentation of the NET region and identification of the NET boundary. **(C)** Skeletonization of the segmented NET and identification of individual filaments. **(D)** Identification and classification of junctions and branch points according to connectivity. **(E)** Distance-transform analysis used to quantify local NET thickness. **(F)** Skeleton-based mesh analysis identifying enclosed pores and their geometric properties. **(G)** Representative local examples of filament thickness, a triple-point junction, and a mesh pore. **(H)** Channel-separated views of B-DNA, Z-DNA, and the merged image. **(I)** Definitions, biological interpretation, and quantitative measurement of extracted NET architectural features. **(J)** Overview of the computational workflow from image preprocessing and NET segmentation through skeletonization, feature extraction, and quantitative output. The analysis pipeline is implemented in the publicly available net-dna-structure-analysis repository. GitHub repository

**Supplementary Figure 2**

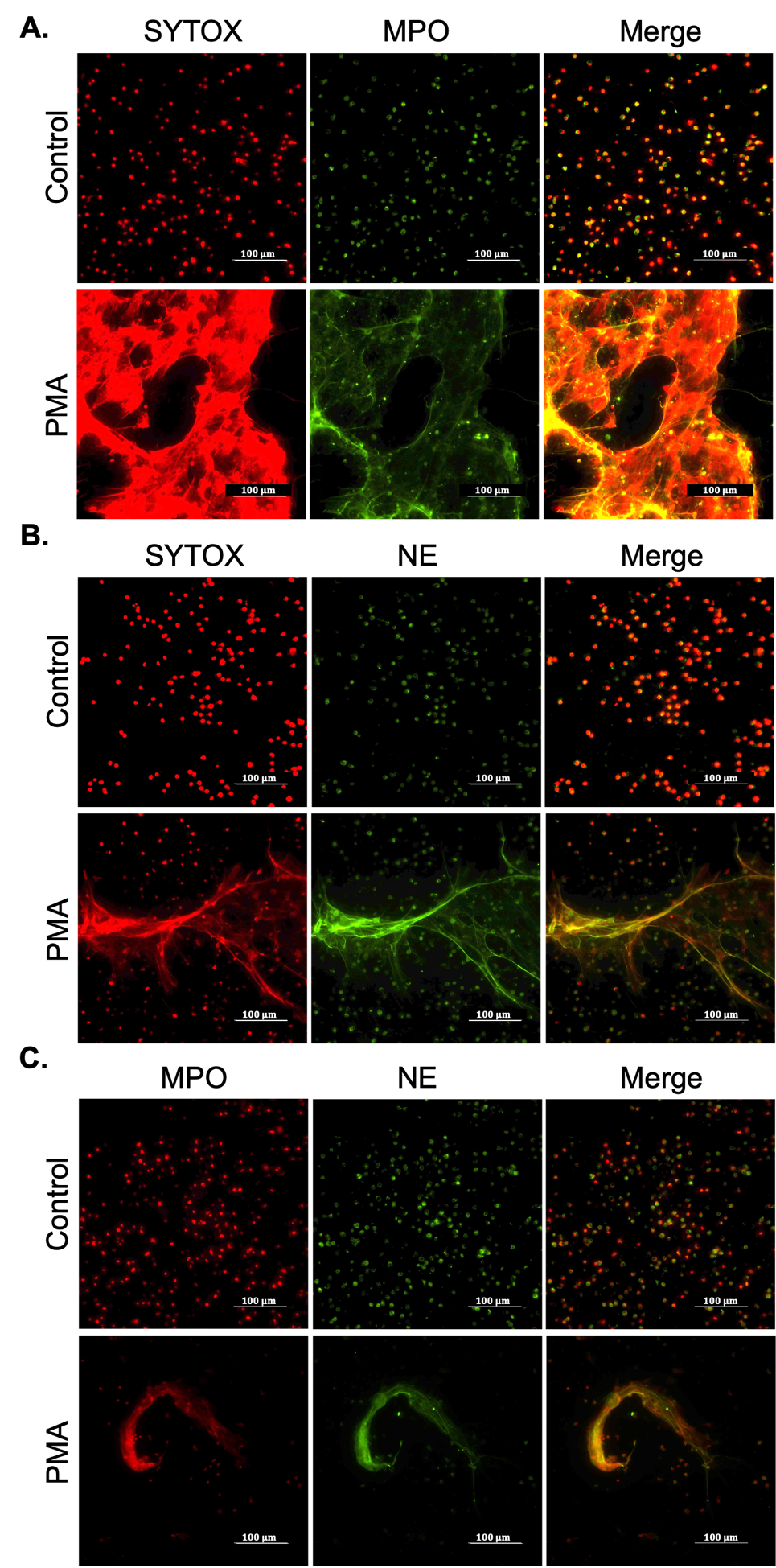

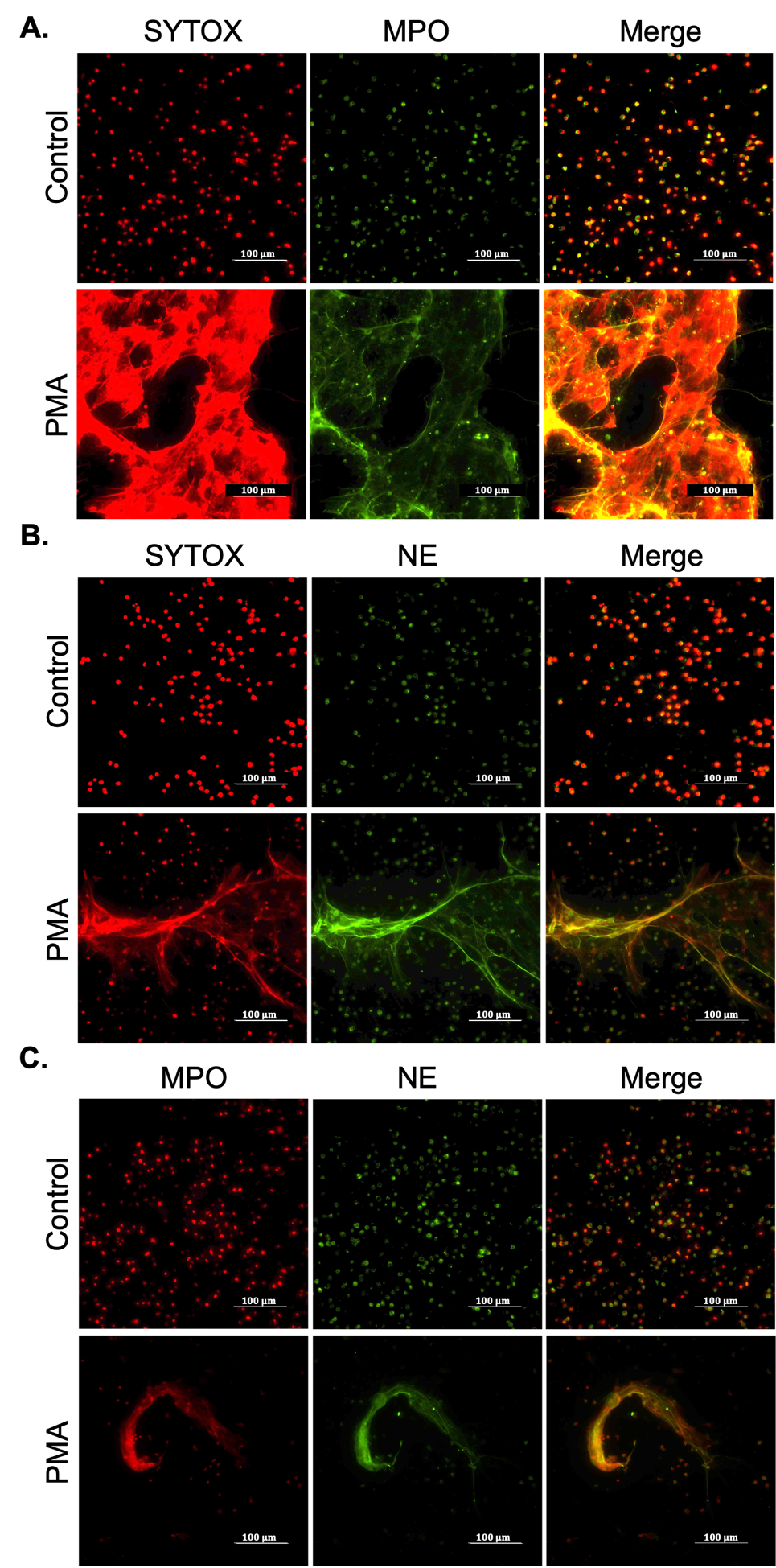

**Supplementary Figure S2: PMA-induced NETs are positive for NETosis markers; myeloperoxidase (MPO) and neutrophil elastase (NE).** Representative immunofluorescence images of unstimulated (Control) and PMA-stimulated (1 µM, 4 h) neutrophils stained for canonical NET markers: **(A)** SYTOX (red) and MPO (green), **(B)** SYTOX (red) and NE (green), and **(C)** MPO (red) and NE (green) co-localization. Scale bars, 100 µm.

**Supplementary Figure 3**

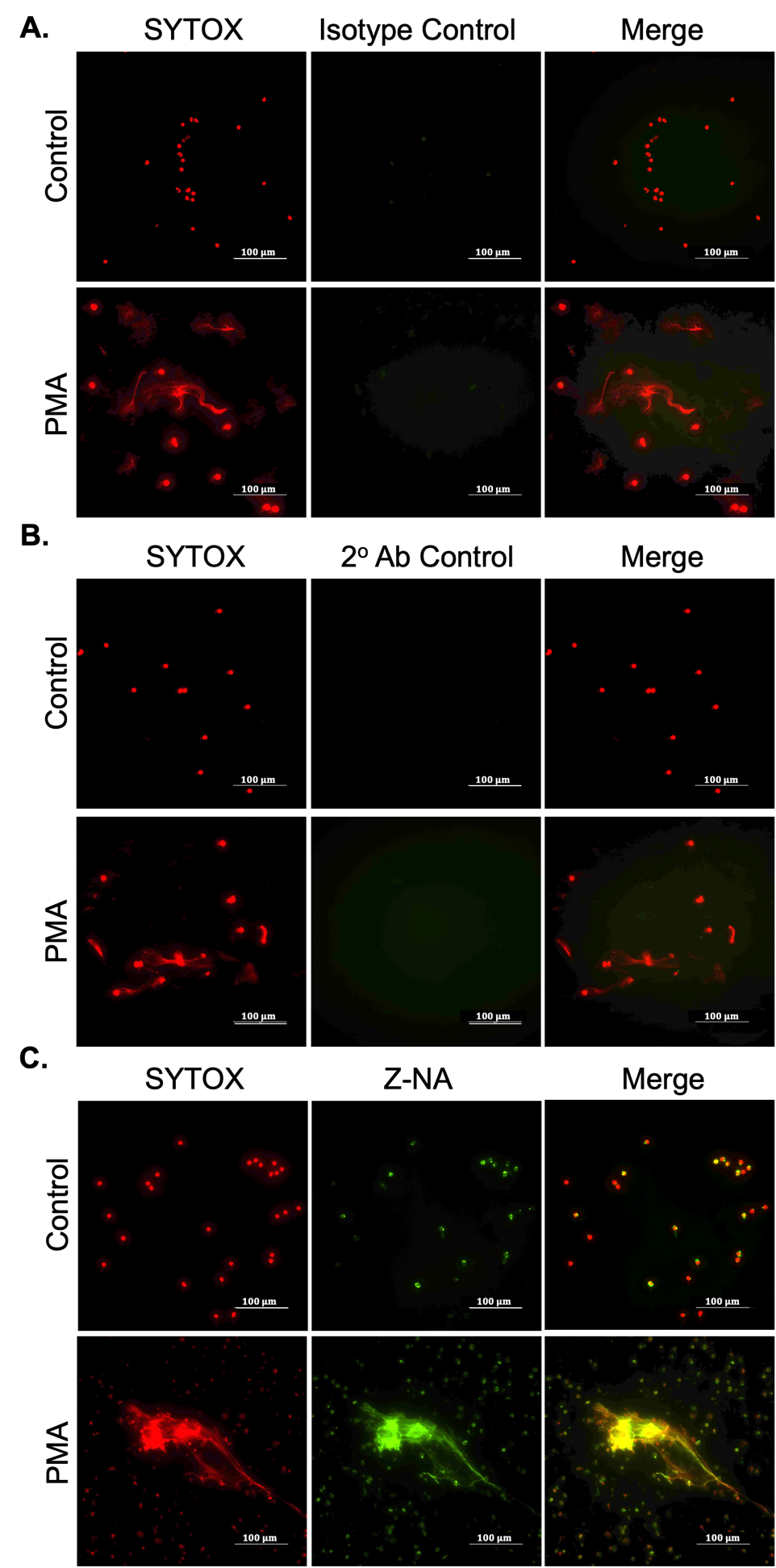

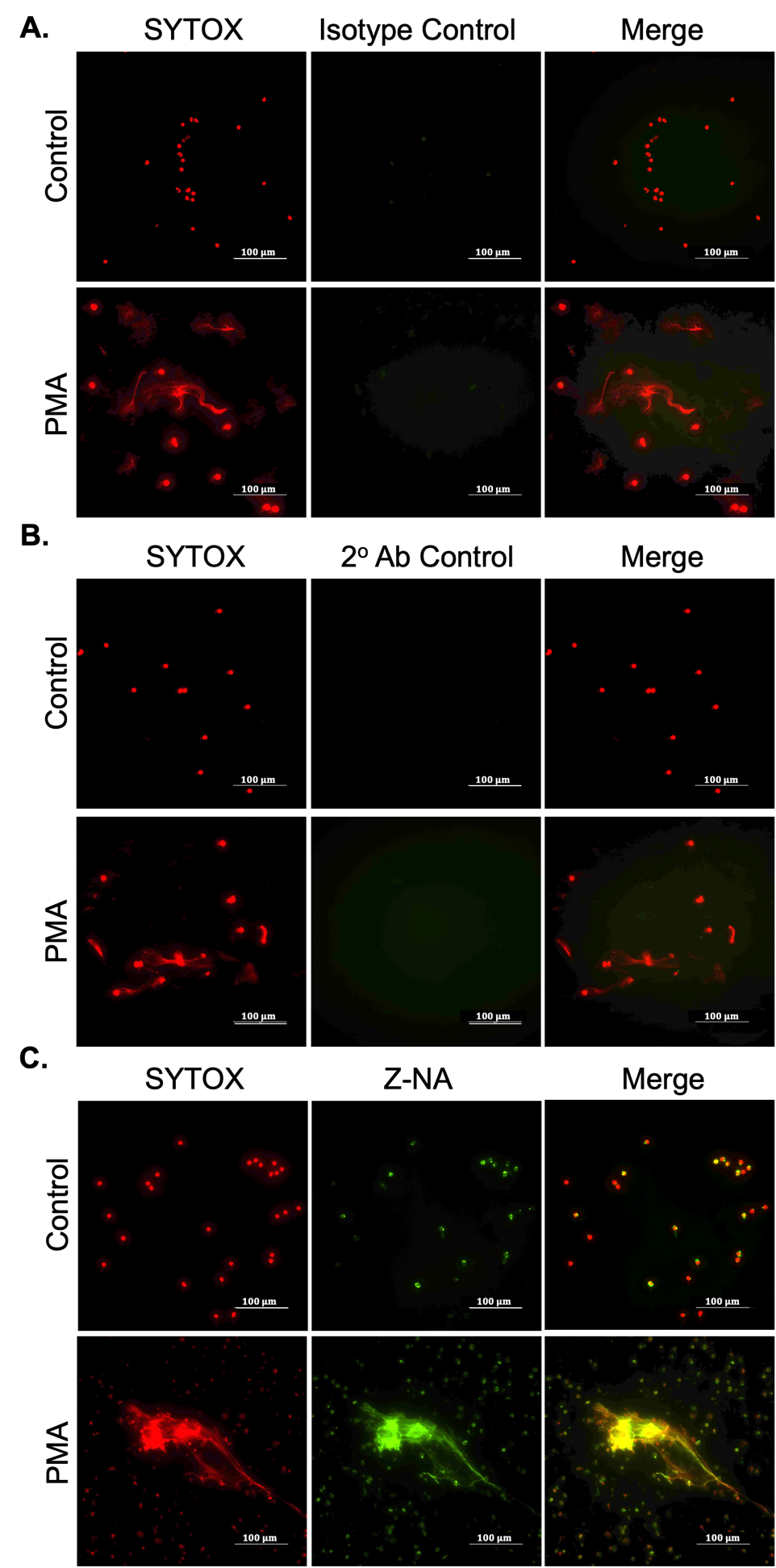

**Supplementary Figure S3: Z22 antibody specificity controls for Z-NA detection in PMA-induced NETs.** Representative immunofluorescence images of unstimulated (Control) and PMA-stimulated (1 µM, 4 h) neutrophils under three antibody conditions: **(A)** isotype-matched IgG control, **(B)** secondary antibody-only control (no primary antibody), and **(C)** Z22 anti–Z-NA antibody (green), non-Z-DNA was stained with SYTOX (red). Scale bars, 100 µm.

**Supplementary Figure 4**

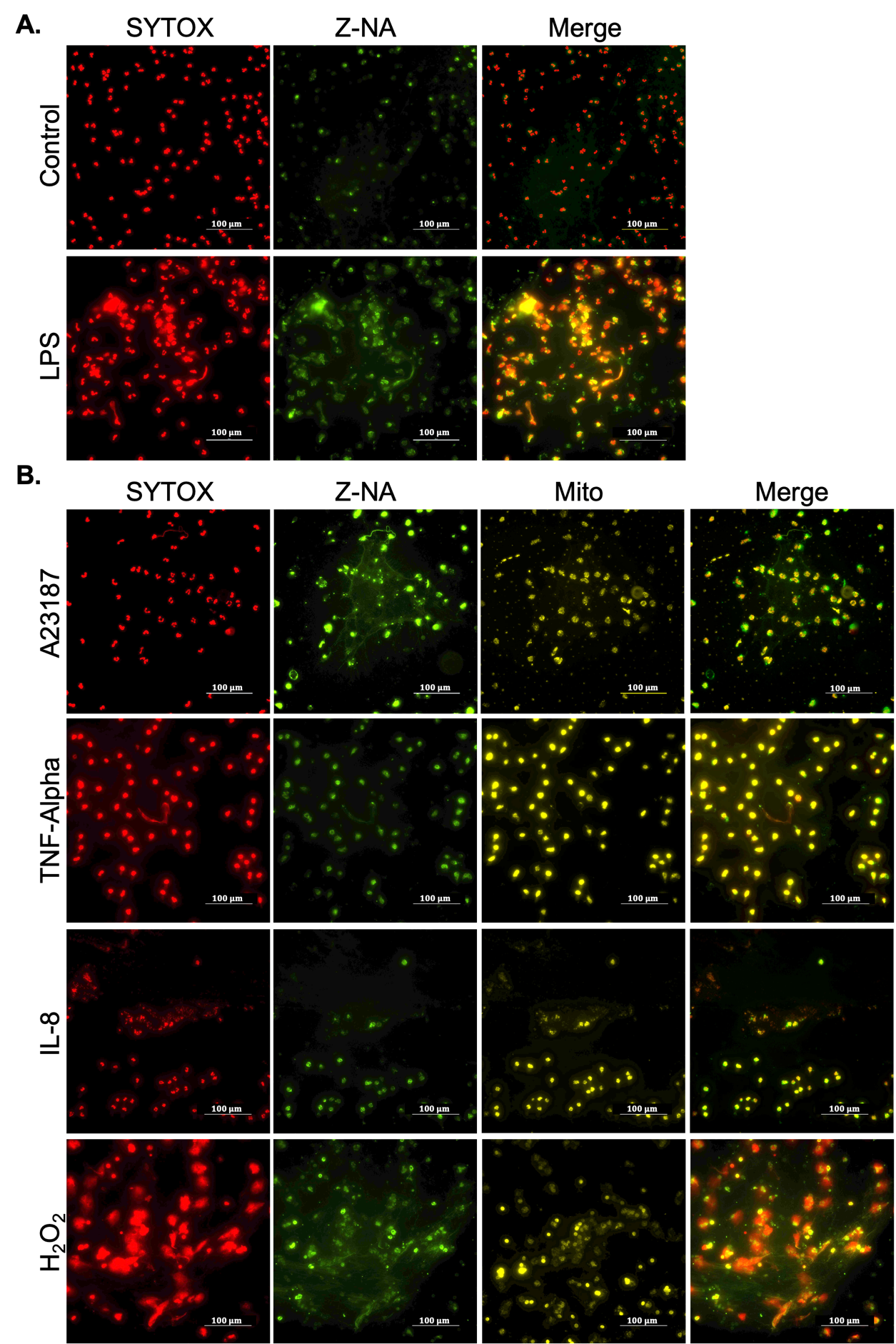

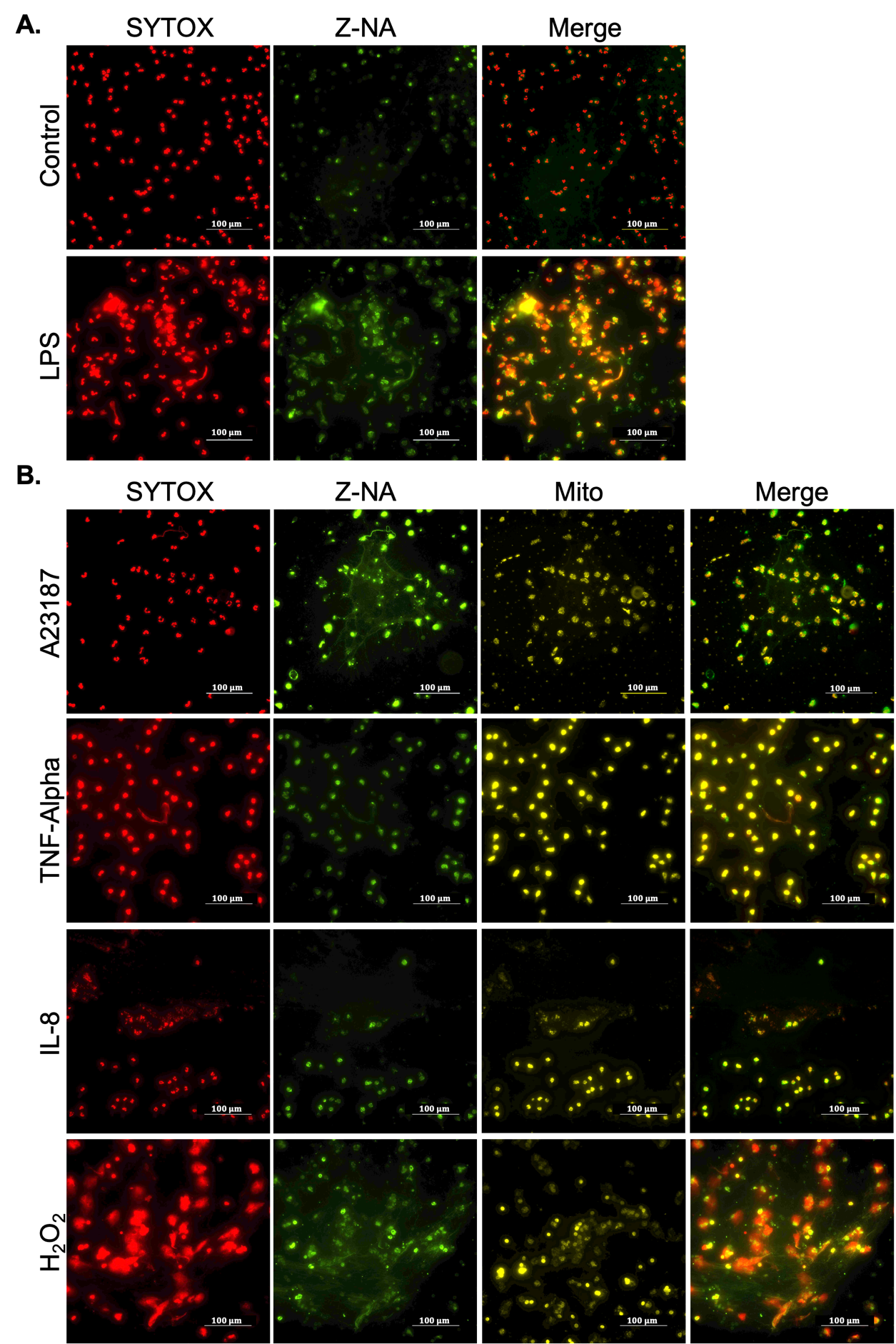

**Supplementary Figure S4: Z-DNA formation across multiple NETosis-inducing stimuli.** Representative immunofluorescence images of neutrophils under the indicated conditions: **(A)** unstimulated control (no treatment) and LPS (100 µg/mL, 4 h); **(B)** A23187 (2 µM, 1 h), TNF-α (20 ng/mL, 4 h), IL-8 (150 ng/mL, 4 h), and H₂O₂ (0.03%, 3 h). Non-Z-DNA was stained with SYTOX (red), Z-DNA was detected using the Z22 anti–Z-DNA antibody (green), and mitochondria were labeled with MitoTracker (Yellow). Merged images show the localization of Z-DNA within total DNA structures. These data correspond to the conditions quantified in Figure 1B. Scale bars, 100 µm.

**Supplementary Figure 5**

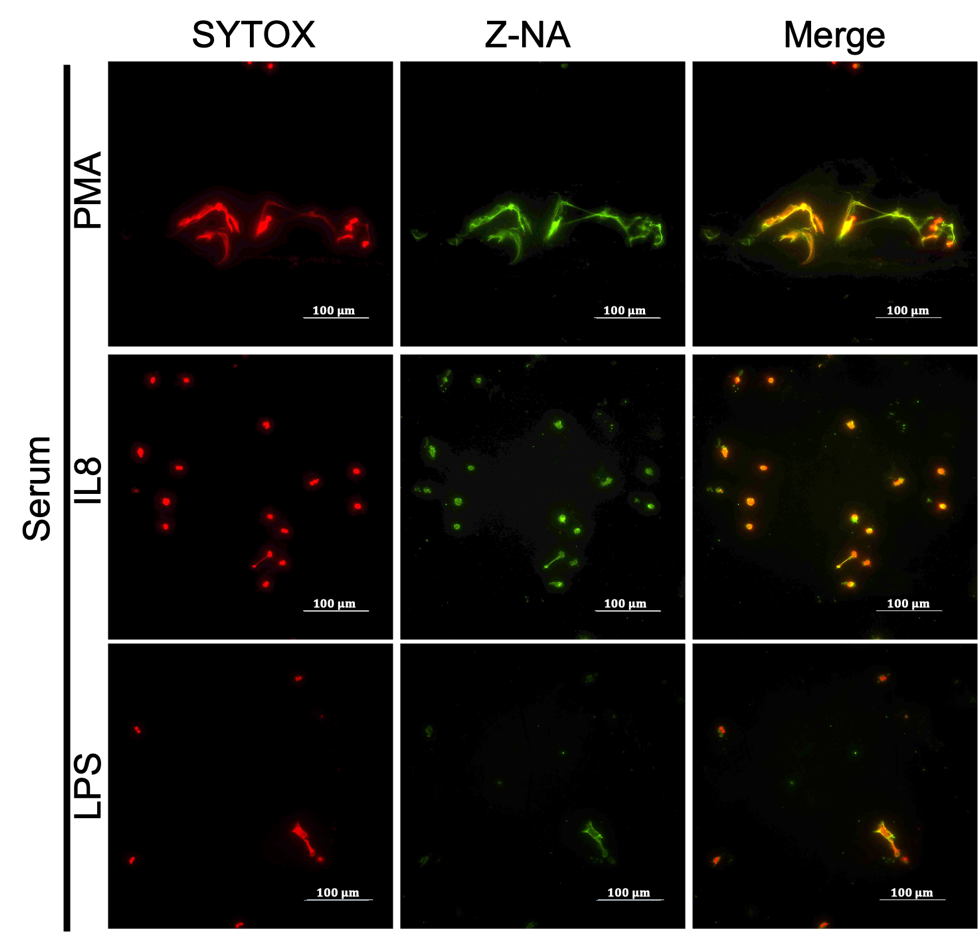

**Supplementary Figure S5:** ***ex vivo* stimulation of NETs results in Z-DNA formation corroborating *in vitro* results.** Representative immunofluorescence images of NETs generated by stimulation with PMA, IL-8, and LPS in the presence of autologous serum. Non-Z-DNA was stained with SYTOX (red), and Z-DNA was detected using the Z22 anti–Z-DNA antibody (green). Merged images show the localization of Z-DNA within total DNA structures. Scale bars, 100 µm.

**Supplementary Figure 6**

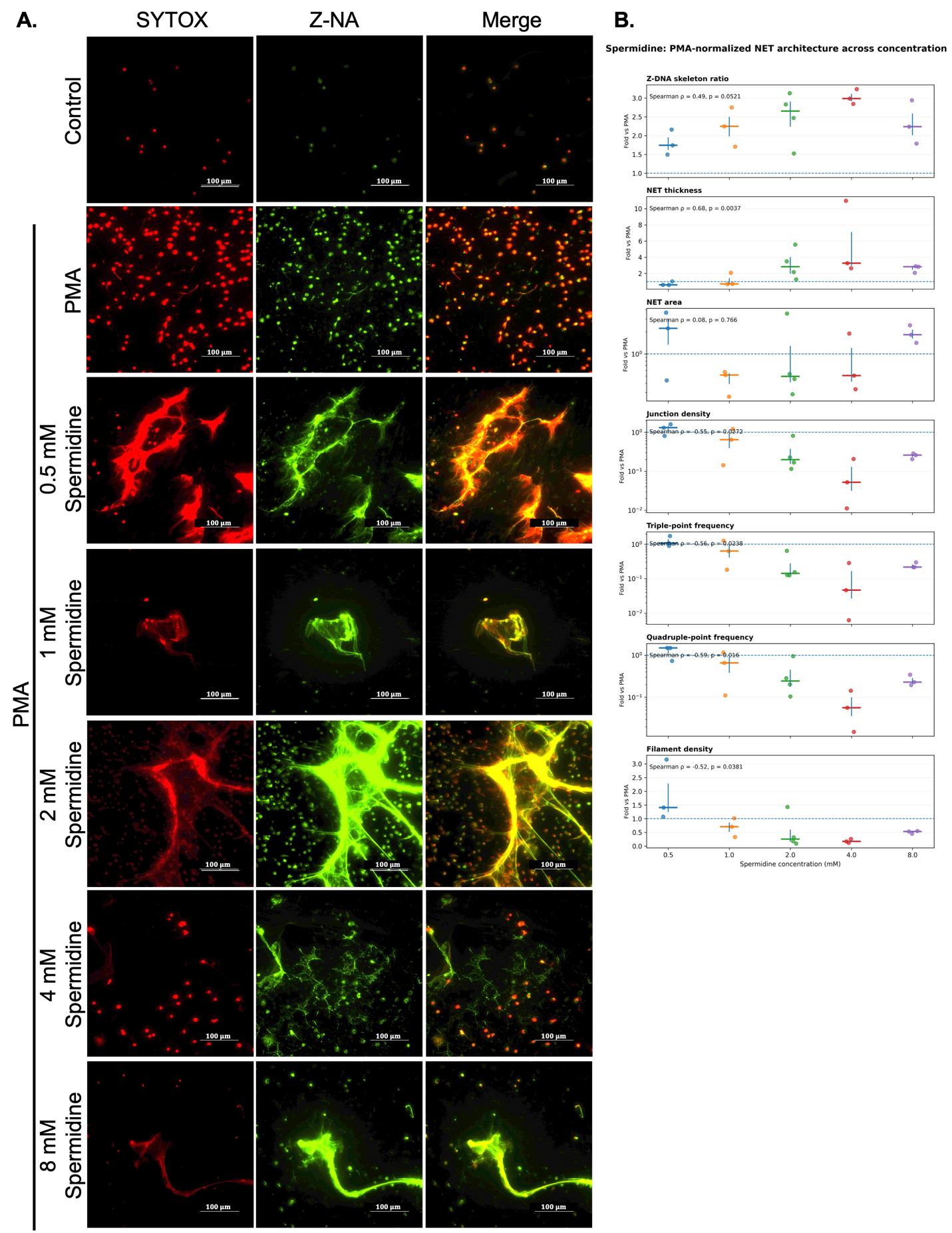

**Supplementary Figure S6: Spermidine promotes Z-DNA enrichment in PMA-induced NETs. (A)** Representative immunofluorescence images of unstimulated control neutrophils and PMA-stimulated neutrophils treated with spermidine trihydrochloride at the indicated concentrations during the final 1 h of stimulation. Z22 anti–Z-NA antibody (green), non-Z-DNA was stained with SYTOX (red). Scale bars, 100 µm. **(B)** Quantitative analysis of spermidine-induced changes in NET architecture, shown from top to bottom as Z-DNA skeleton ratio, NET thickness, NET area, junction density, triple-point frequency, quadruple-point frequency, and filament density. Values are expressed as fold change relative to the corresponding PMA-only NETs. Individual points represent images, with larger symbols and error bars indicating mean ± SD. Horizontal dashed lines indicate the PMA reference value (1.0-fold). Spearman correlation coefficients and p values are shown within each panel.

**Supplementary Figure 7**

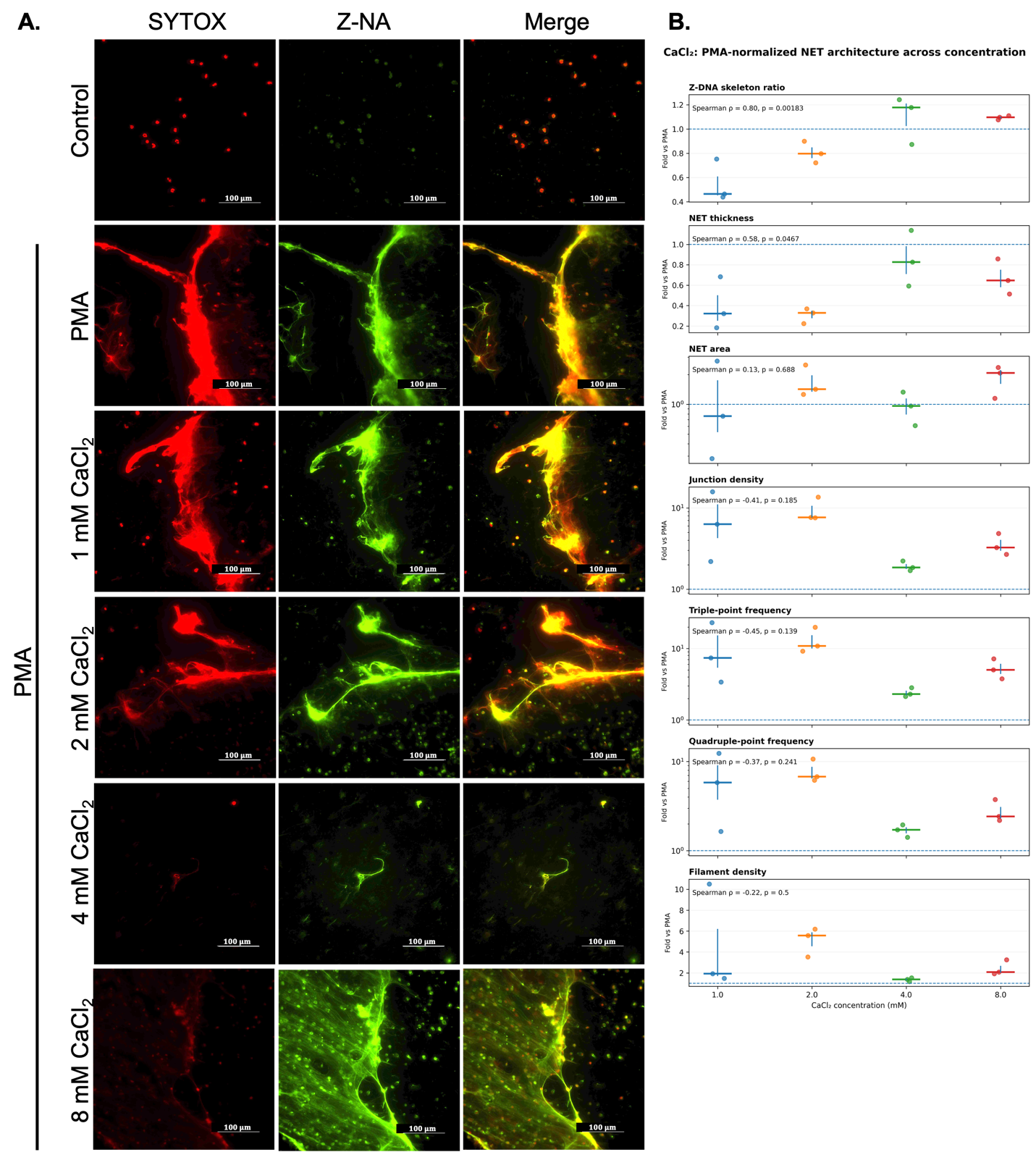

**Supplementary Figure S7: CaCl2 promotes Z-DNA enrichment in PMA-induced NETs.** Representative immunofluorescence images of unstimulated control neutrophils and PMA-stimulated neutrophils treated with CaCl2 at the indicated concentrations during the final 1 h of stimulation. Z22 anti–Z-NA antibody (green), non-Z-DNA was stained with SYTOX (red). Scale bars, 100 µm. (B) Quantitative analysis of CaCl2-induced changes in NET architecture, shown from top to bottom as Z-DNA skeleton ratio, NET thickness, NET area, junction density, triple-point frequency, quadruple-point frequency, and filament density. Values are expressed as fold change relative to the corresponding PMA-only NETs. Individual points represent images, with larger symbols and error bars indicating mean ± SD. Horizontal dashed lines indicate the PMA reference value (1.0-fold). Spearman correlation coefficients and P values are shown within each panel.

**Supplementary Figure 8**

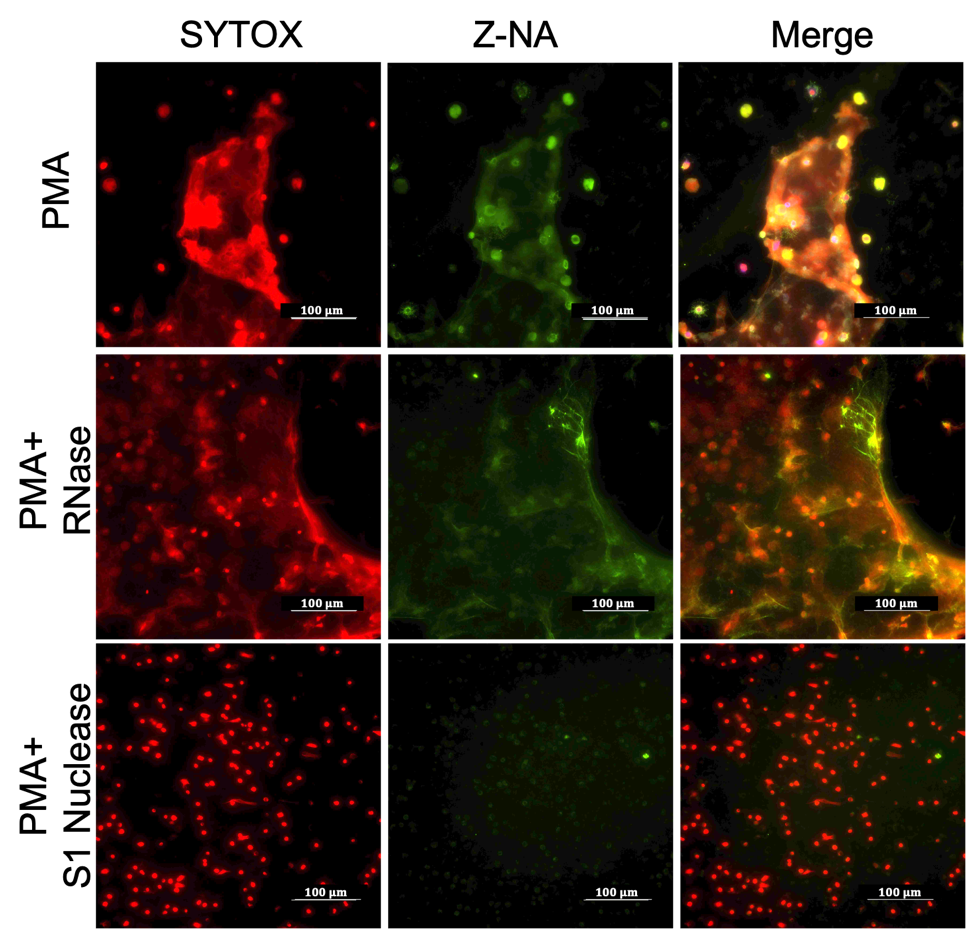

**Supplementary Figure S8:** Representative immunofluorescence images of NETs generated by PMA stimulation (1 µM, 4 h) and subsequently treated with RNase or S1 nuclease. Non-Z-DNA was stained with SYTOX (red), and Z-DNA was detected using the Z22 anti–Z-DNA antibody (green). Merged images show the localization of Z-DNA within total DNA structures. Scale bars, 100 µm.

**Supplementary Figure 9**

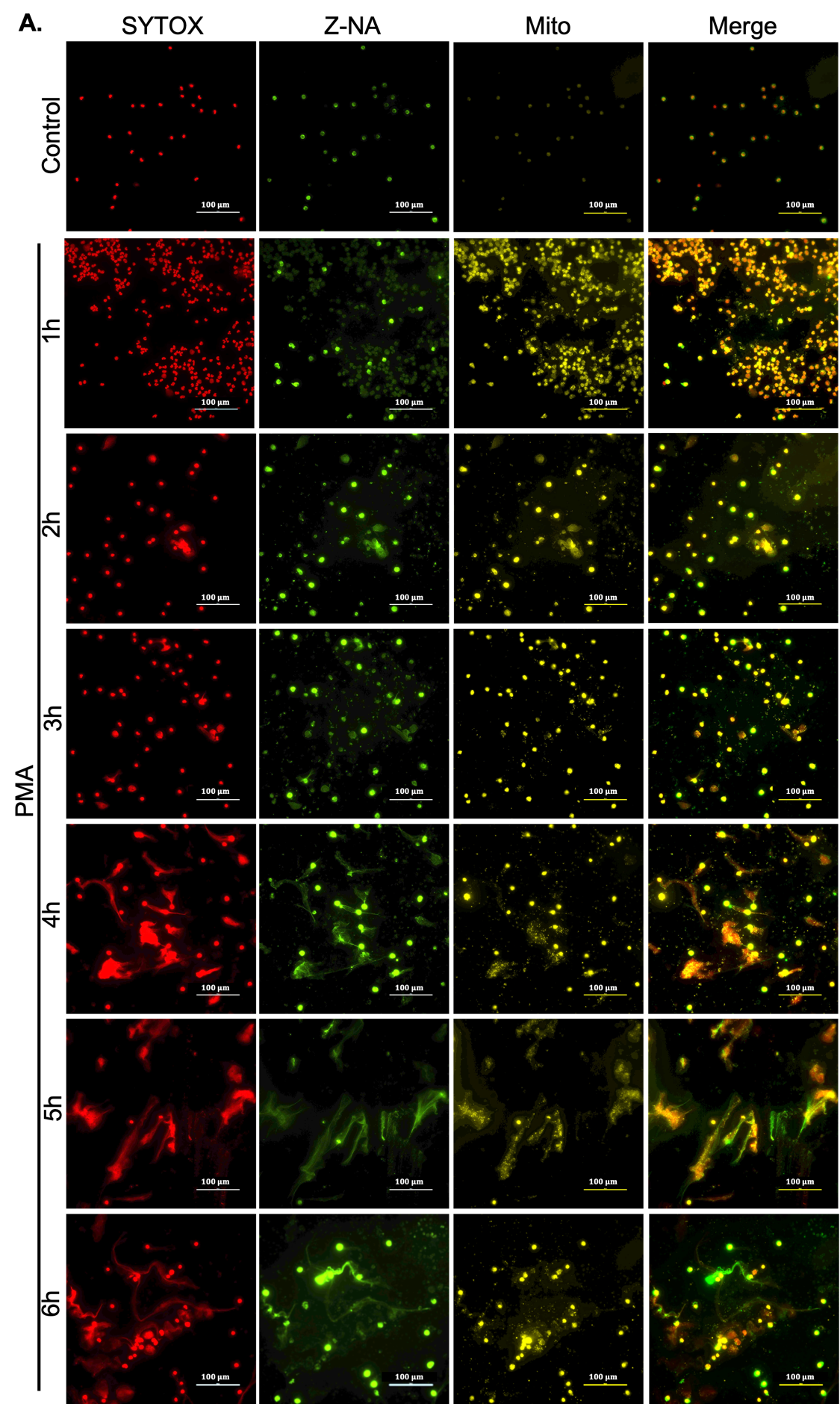

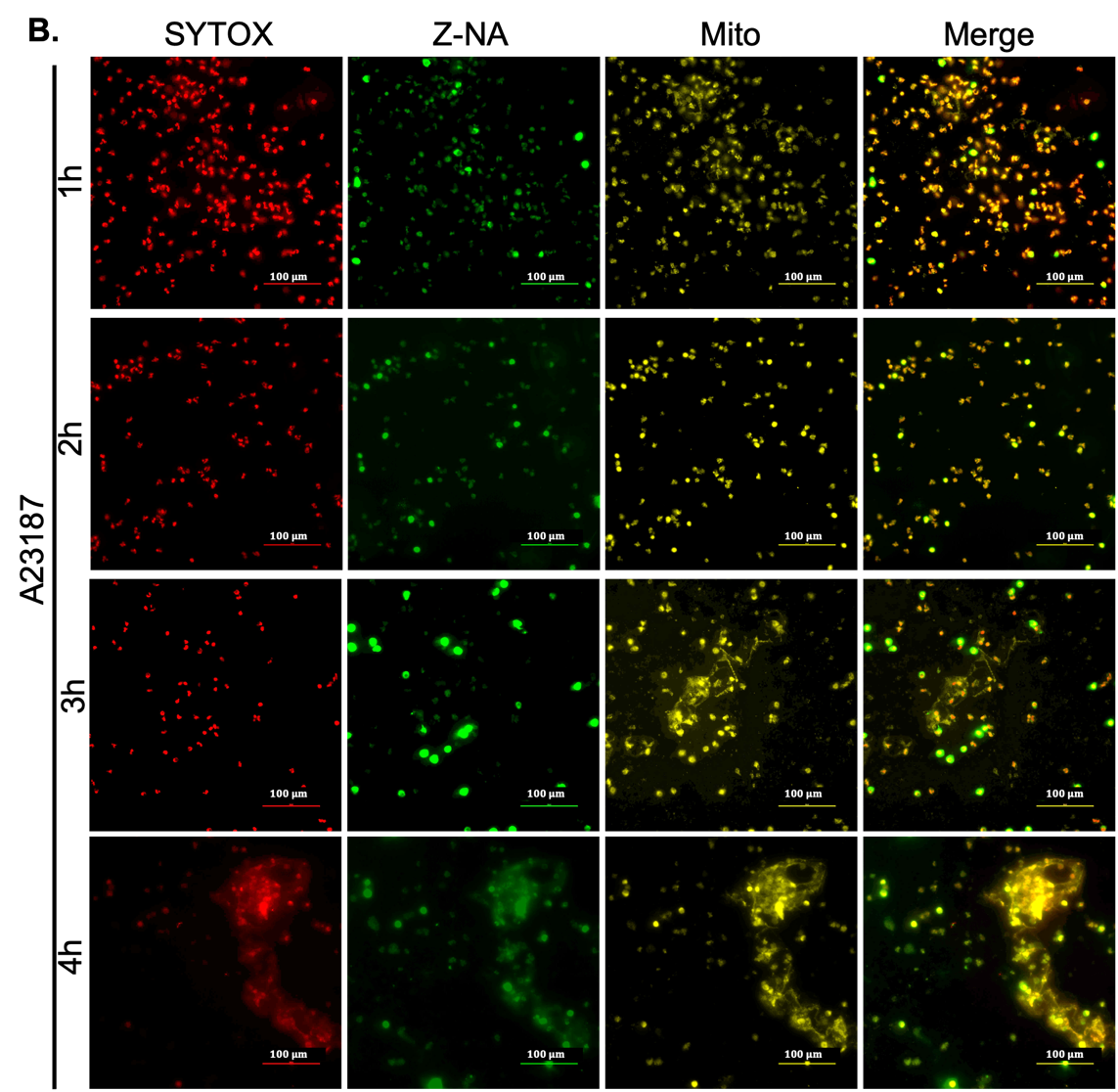

**Supplementary Figure S9:** Representative immunofluorescence images of neutrophils following **(A)** PMA- and **(B)** A23187-induced NETosis at the indicated time points, stained for Z-DNA (Z-NA, green), mitochondria (MitoTracker, yellow), and non-Z-DNA (SYTOX, red), with corresponding merged images. Scale bars, 100 µm.

**Supplementary Figure 10**

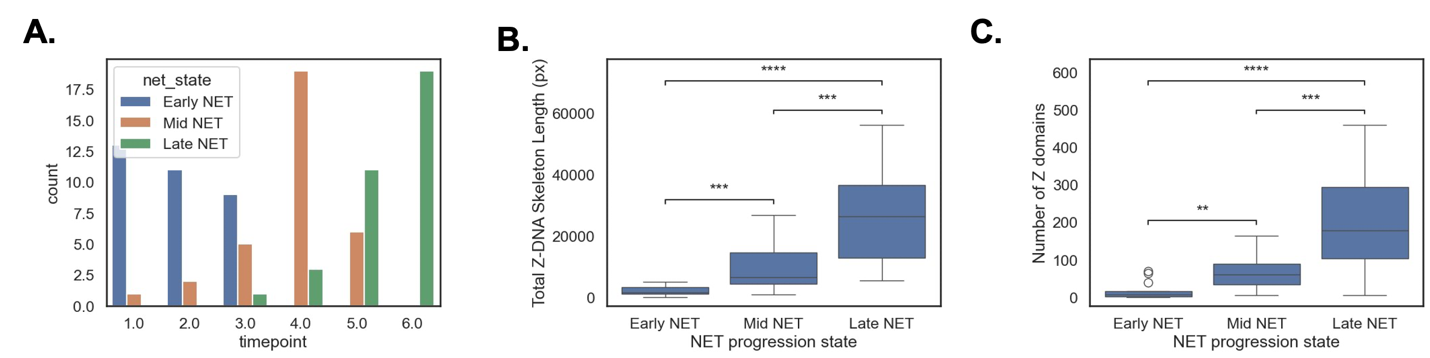

**Supplementary Figure S10:** NET progression state classification accounts for temporal heterogeneity and clearly delineates development of Z-DNA with maturation of NETs. **(A)** classification of replicates and different fields of view into states based on the progression score captures heterogeneity within each time point. **(B)** and **(C)** Total Z-DNA skeleton length and number of Z domains significantly increase with maturation of NETs.

**Supplementary Figure 11**

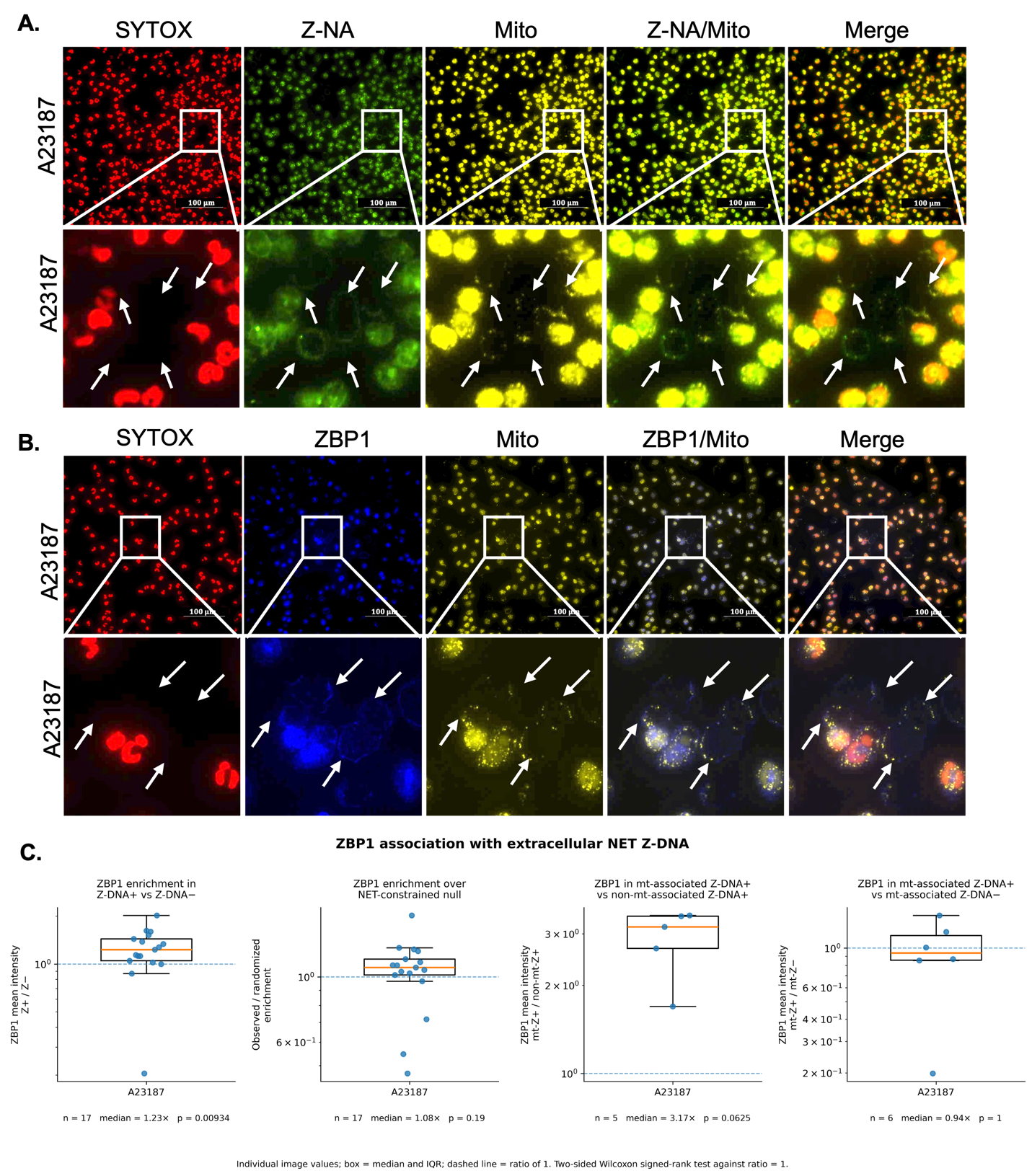

**Supplementary Figure S11: (A) Z-DNA colocalize with mitochondrial compartments and (B) ZBP1 in extracellular NET structures.** Representative immunofluorescence images of neutrophils stimulated with A23187 (A23, 2 µM, 1 h) to induce mitochondrial DNA release. non-Z-DNA was stained with SYTOX (red), Z-DNA was detected using the Z22 anti–Z-DNA antibody (green), mitochondria were labeled using MitoTracker (yellow), and ZBP1 was detected using an anti-ZBP1 (blue) antibody. Insets (arrows) represent magnified views of the corresponding regions of interest (ROIs) from the panels above. Scale bars, 100 µm. **(C)** ZBP1 association with extracellular Z-DNA was assessed in A23187-induced NETs by comparing ZBP1 signal within Z-DNA-positive (Z-DNA⁺) regions with Z-DNA-negative (Z-DNA⁻) regions and with a NET-constrained randomized null distribution. From left to right: ZBP1 mean intensity was enriched in Z-DNA⁺ relative to Z-DNA⁻ regions (n = 17 images; median enrichment = 1.23-fold; P = 0.00934); observed ZBP1 enrichment relative to the NET-constrained randomized null was modest and did not reach significance (n = 17; median = 1.08-fold; P = 0.19); ZBP1 was further enriched in mitochondria-associated Z-DNA⁺ regions relative to non-mitochondria-associated Z-DNA⁺ regions (n = 5; median enrichment = 3.17-fold; P = 0.0625); and ZBP1 signal in mitochondria-associated Z-DNA⁺ regions did not differ from that in mitochondria-associated Z-DNA⁻ regions (n = 6; median enrichment = 0.94-fold; P = 1). Box plots show median and IQR, with whiskers indicating the observed range; individual points represent images. Dashed horizontal lines indicate an enrichment ratio of 1.0. Two-sided Wilcoxon signed-rank tests were used for comparisons against a ratio of 1.0.

**Supplementary Figure 12**

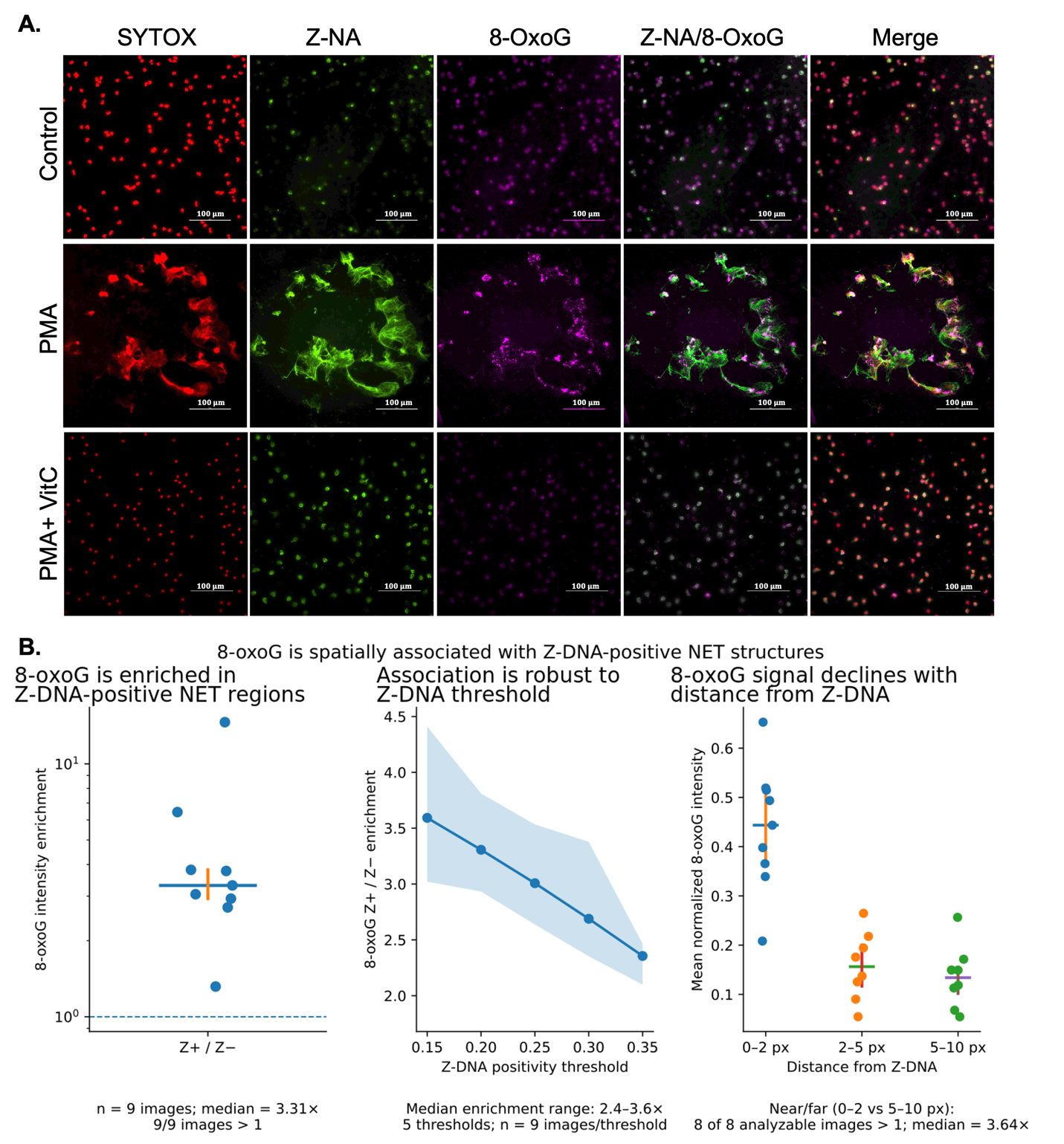

**Supplementary Figure S12: (A)** Asymmetric colocalization between 8-oxoG and Z-DNA in NETs. Representative immunofluorescence images of neutrophils under unstimulated conditions and following induction of NETosis with PMA (1 µM, 4 h), with or without vitamin C (VitC, 2 mM, 30 min) treatment. non-Z-DNA was stained with SYTOX (red), Z-DNA was detected using the Z22 anti–Z-DNA antibody (green), and oxidative DNA damage was visualized using an anti–8-oxoG antibody (magenta). Merged images show spatial co-localization of Z-DNA with 8-oxoG–positive regions within extracellular DNA structures. **(B)** Spatial association of 8-oxoG with Z-DNA-positive NET structures. Left - 8-oxoG intensity was enriched within Z-DNA-positive (Z⁺) NET regions relative to Z-DNA-negative (Z⁻) regions, with a median enrichment of 3.31-fold across 9 images; 8 of 9 images showed enrichment greater than 1 (dashed line). Center - The association between 8-oxoG and Z-DNA remained robust across Z-DNA positivity thresholds from 0.15 to 0.35, with median enrichment values ranging from 2.4- to 3.6-fold (n = 9 images). Right - 8-oxoG signal decreased with increasing distance from Z-DNA. In 8 of 8 analyzable images, 8-oxoG intensity in the 0–2-pixel region immediately proximal to Z-DNA exceeded that in the 5–10-pixel region, with a median near-to-far enrichment of 3.64-fold. Individual points represent images; horizontal lines indicate group means.
